# Myoglobin-derived iron causes wound enlargement and impaired regeneration in pressure injuries of muscle

**DOI:** 10.1101/2022.03.07.483146

**Authors:** N. Jannah M. Nasir, Hans Heemskerk, Julia Jenkins, N. Hidayah Hamadee, Ralph Bunte, Lisa Tucker-Kellogg

**Author notes:** Prof. Tucker-Kellogg, 8 College Road, Singapore 169857.

## Abstract

The reasons for poor healing of pressure injuries are poorly understood. Vascular ulcers are worsened by extracellular release of hemoglobin, so we examined the impact of myoglobin (Mb) iron in murine muscle pressure injuries (mPI). Tests used Mb-knockout or treatment with deferoxamine iron chelator (DFO).

Unlike acute injuries from cardiotoxin, mPI regenerated poorly with a lack of viable immune cells, persistence of dead tissue (necro-slough), and abnormal deposition of iron. However, Mb-knockout or DFO-treated mPI displayed a reversal of the pathology: decreased tissue death, decreased iron deposition, decrease in markers of oxidative damage, and higher numbers of intact immune cells. Subsequently, DFO treatment improved myofiber regeneration and morphology.

We conclude that myoglobin iron contributes to tissue death in mPI. Remarkably, a large fraction of muscle death in untreated mPI occurred later than, and was preventable by, DFO treatment, even though treatment started 12 hours after pressure was removed. This demonstrates an opportunity for post-pressure prevention to salvage tissue viability.

## Introduction

Pressure injuries (also called pressure ulcers, bedsores, decubiti, or pressure sores) are tissue damage caused by sustained pressure. They are extremely painful (1), costly to prevent and treat (2), and they increase the risk of patient sepsis and death (3). Tissue death can be caused by mechanical deformation, ischemia, or both. Ischemia is often studied as ischemia-reperfusion injury, but poor clearance of damage factors may have unappreciated importance (4).

Pressure injuries often heal poorly (5), especially if they involve deeper layers such as muscle (6, 7), but the reasons are for poor quality and quantity of healing are not clear. Some cases can be explained by complications or co-morbidities (infection, incontinence, poor circulation, hyperglycemia, chronic re-injury, advanced age) but pressure injuries can affect any immobile person (e.g., young adults with spinal cord injury). In this study, we ask whether some aspect of pressure-induced injury is intrinsically inhospitable to regeneration and in need of intervention.

Chronic ulcers of veins or arteries (e.g., venous stasis ulcers, sickle cell ulcers), have high levels of extracellular hemoglobin (Hb) released in the wound. For example, many have deposits of hemosiderin. Extracellular Hb and its breakdown products (e.g., hemin, iron) create oxidative stress (8, 9) and other effects that are detrimental to regeneration. For example, Hb decreases nitric oxide for angiogenesis (10, 11), and signals as a DAMP (damage-associated molecular pattern) to increase inflammation (12, 13). Systems exist to detoxify Hb (14), but Hb is also an innate immune factor with evolutionarily-conserved antimicrobial function. When extracellular Hb is activated by proteolytic cleavage, bacterial binding, or conformational change, it increases production of ROS (reactive oxygen species) (15); this process has been called “oxidative burst without phagocytes” (15, 16). Our earlier work used evolutionary conservation to identify ROS-producing fragments of Hb and crosstalk with tissue factor coagulation (17, 18). Therefore, globin proteins have multiple functions that may be detrimental to chronic wounds (19).

Myoglobin release into plasma or urine has been observed after muscle pressure injuries (mPI) in multiple studies including DTI (deep tissue injury) (20–24), but was studied as a readout of damage rather than a source of damage. We reason by analogy to hemoglobin that extracellular myoglobin might create a hostile wound environment. An extracellular environment oxidizes globins to a ferric (Fe^3+^) state, which can be further oxidized to ferryl (Fe^IV^ = O) globin in the presence of endogenous peroxides such as hydrogen peroxide (25). Hydrogen peroxide is ubiquitous in contexts of cell stress, mitochondrial permeabilization, and cell death (26). Ferryl-Mb can oxidize macromolecules directly (8, 27, 28) and can form heme-to-protein cross-links (29). Most importantly, ferryl myoglobin can participate in a catalytic cycle of pseudo-peroxidase activity (redox cycling) (30). In a tissue context, myoglobin can induce ferroptosis, which is a form of non-apoptotic cell death associated with iron and characterized by lipid peroxidation (31). Dissociation of myoglobin into free heme or iron results in additional forms of toxicity, as described for hemoglobin.

We hypothesize that mPI will have Mb-dependent pathologies, and that introducing Mb-knockout or iron chelation therapy will partially normalize the mPI pathologies. Deferoxamine (DFO), also known as desferrioxamine or desferoxamine, is an FDA-approved small-molecule iron chelator that improves iron overload syndromes (32, 33). DFO binds free iron and heme at a 1-to-1 ratio (32), scavenges free radicals (25, 34), reduces ferryl myoglobin to ferric myoglobin (28, 35), inhibits cross-linking of heme to protein (9), and prevents the formation of pro-oxidant globin and heme species. DFO can function as an activator of Hif1α (19, 36), a tool for promoting angiogenesis (37, 38), an antioxidant (39), or can join an anti-ischemic cocktail (40). DFO appears in hundreds of studies of ischemic or inflammatory pathologies. In our study, subcutaneous DFO is used for testing the hypothesized role of myoglobin iron and as an anti-DAMP therapy for combating local iron overload.

Assessing the contribution of myoglobin iron to pressure injury pathophysiology provides an opportunity to test several additional hypotheses about pressure injuries. First, our prior work in mathematical modeling (41) predicted that oxidative stress from myoglobin and other DAMPs could create ***secondary progression*** of pressure ulcers. Secondary progression means that otherwise viable tissue dies later from the environmental consequences of injury, rather than dying directly from the original injury. Pressure injuries are known to have gradual expansion of tissue death (42), consistent with secondary progression, but blocking secondary progression has not been clinically recognized as a goal for intervention (43). Therefore, our studies are designed to test whether tissue margins can be saved from dying, if we initiate iron chelation therapy 12 hours after pressure has ended. Second, we hypothesize that iron chelation therapy, by improving the early stages of injury response, will lead to better muscle tissue architecture (better morphogenesis) in long-term regeneration, even after treatment has ended. This hypothesis will be tested by breeding inducible fluorescence into satellite cells (muscle stem cells bearing Pax7); this fluorescence causes newly regenerated muscle fibers light up against the dark background of pre-existing muscle. Third, establishing a pressure injury mouse model provides an opportunity to learn how much of the poor healing is independent of co-morbidities and complications. To the best of our knowledge, pressure injuries have never been assessed for poor regeneration under aseptic conditions in young, healthy animals. We hypothesize that even under these ideal circumstances, mPI will heal slowly and incompletely. Our fourth and final additional hypothesis is inspired by prior studies of blood-related conditions in which high levels of hemoglobin or heme/hemin could impair the survival, chemotaxis, and phagocytosis (11, 44–50) of phagocytic cells. Given that pressure ulcers often have slough or eschar, we hypothesize that necrotic tissue will persist in the mPI wound bed, and that sterile mPI will have slough, despite the absence of bacterial biofilm. If correct, this would imply that slough by itself is not sufficient to indicate infection (or bacterial colonization) of a wound.

## Results

### Magnet-induced pressure injury causes delayed healing and failure of muscle regeneration

To compare wound healing between acute and chronic wounds, we injured the dorsal skinfold of mice using either cardiotoxin (CTX) or pressure (Suppl. Fig 1A-B) in healthy young adult mice under specific pathogen-free conditions. Both groups of mice received sham-treatment (injected with 0.9% saline subcutaneously for 16 days, or until mouse sacrifice, whichever was sooner). The normal uninjured mouse skinfold contains the following parallel layers: a thin epithelium (epidermis), a thicker layer of dermis, dermal white adipose tissue (dWAT), a muscle layer called the panniculus carnosus (PC), and a wavy layer of loose areolar tissue (Suppl. Fig 1C). Despite comparable diameters of dead muscle between CTX and mPI at day 3, the wound diameters were vastly different at day 10 (Suppl. Table 1). In the CTX injury at day 3, many blood vessels were intact and carrying red blood cells, but in mPI at day 3, intact vasculature was not observed in the compressed region (Suppl. Fig 2). Prior work showed that pressure affects blood and lymphatic vessels in many ways including potentially loss of flow (4, 51, 52). After CTX killed the panniculus muscle, substantial muscle regeneration occurred by 10 days, in which immature muscle fibers displayed central nuclei (Fig 1A). At 40 days after acute injury, muscle was completely regenerated and mature (evidenced by peripherally-located nuclei, Fig 1C). In contrast, the pressure-injured wound bed remained filled with dead tissue at day 10 (Fig 1B, 1E-F). Our pressure injuries showed no signs of infection and no epibole (Fig 1E). The dead epidermis, dermis, dWAT and panniculus layers were pushed upward at day 7 ± 2 as slough (necroslough, per the nomenclature of (53)) and remained at the surface, eventually becoming a dry eschar (Fig 1E). When the eschar dropped off (by day 15), the size of the epithelial opening was smaller than the eschar, meaning that re-epithelialization had occurred underneath the eschar. Re-epithelialization completed at day 21 ± 2. Despite successful closure of the epithelial layer, pressure injuries at 40 days had only partial regeneration of the panniculus carnosus layer (Fig 1D, 1G). At 90 days after pressure injury, the dermis and epidermis had regenerated, but a hole remained in the panniculus muscle layer (Fig 1H), indicating a failure to regenerate (54). Suppl. Fig 3 summarizes the timelines. We conclude that the mPI healed poorly.

**Figure 1:**
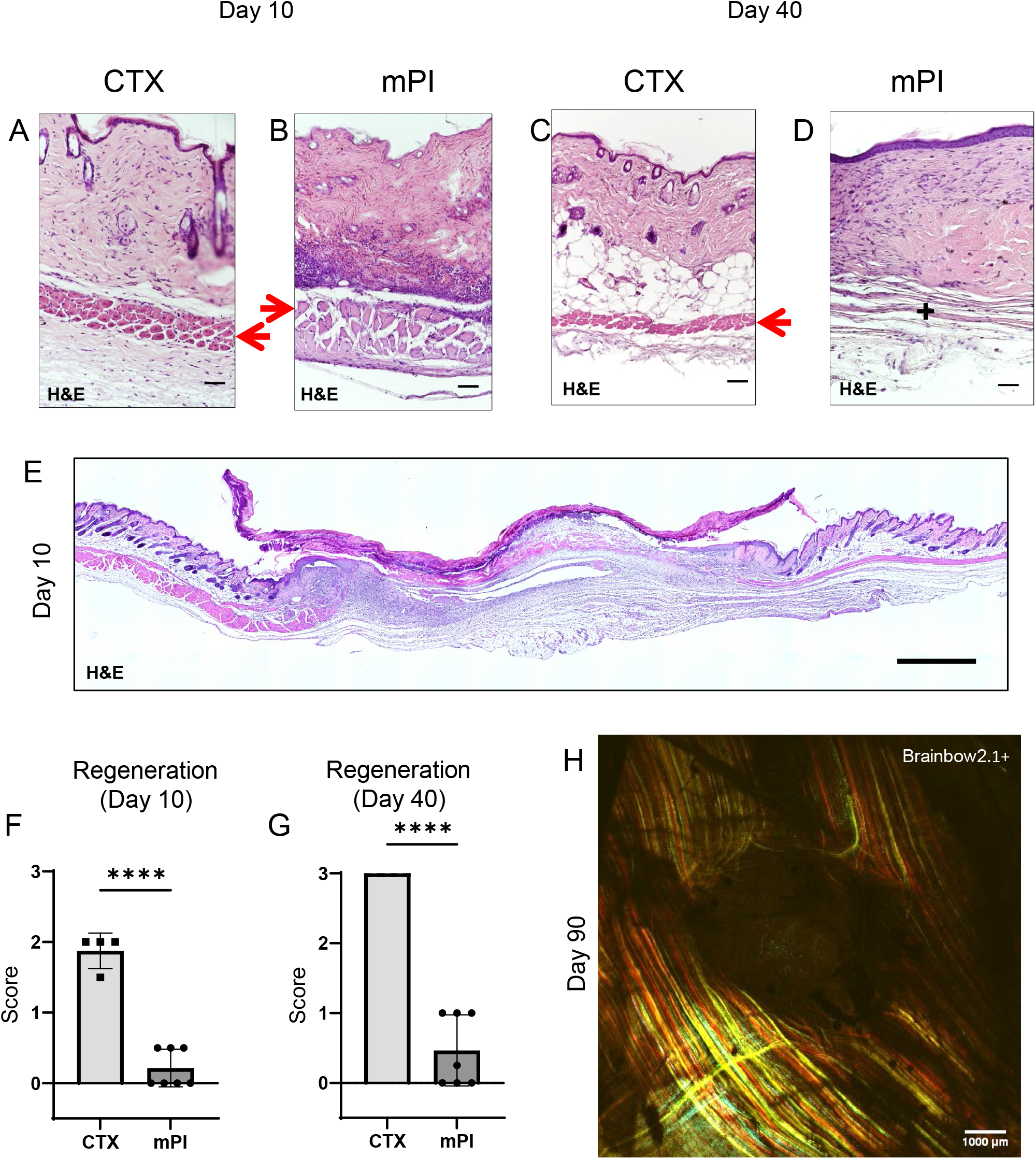
Poor regeneration of muscle pressure injury (mPI), a model of chronic wound, compared with cardiotoxin (CTX), a model of acute injury. H&E-stained sections of saline-treated wound tissues (A-B) at day 10 post-injury and (C-D) day 40 post-injury. Red arrows point to the panniculus carnosus layer. ‘+’ indicates where the panniculus layer should be. Scale bars: 50 μm. Uninjured control tissue, stained with H&E, is shown in Suppl. Fig 1C. (E) Full cross-section of H&E-stained mPI, including uninjured edges, at day 10 post-injury. Note the eschar attached to the wound surface. Scale bar: 500 μm. (F-G) Comparison of the regeneration scores for the panniculus layer between mPI and CTX injuries at day 10 and day 40. (H) A multi-channel confocal image of the panniculus layer shows a round hole remains in the muscle, 90 days after mPI. Scale bar: 1000 μm. All injuries are saline-treated to permit comparison against later mPI treatments.

### Compressed regions of pressure injury display an absence of viable immune cells

To investigate why dead tissue remains at day 10, we studied tissue sections from day 3 post-injury. Both cardiotoxin-induced and pressure-induced injuries had comprehensive death of muscle tissue, as indicated by karyolysis (dissolution of nuclear components), karyorrhexis (fragmentation of the nucleus) and acidification (eosinification) in H&E staining of the panniculus carnosus (Fig 2B). Cardiotoxin and pressure injuries had a difference in morphology: cells in pressure-injured tissues were flattened, and the thickness of the panniculus muscle layer was half of uninjured (Fig 2A-C; p < 0.0001). Even more striking was the difference in immune cell numbers. The muscle layer of cardiotoxin wounds had six-fold higher levels of immune cell infiltrate than mPI (p < 0.0001; Fig 2A-B, and Fig 2D). The panniculus layer of mPI was nearly devoid of intact immune cells. Some viable immune cells were found at the margins of the wound (at the boundary between injured and uninjured tissue), but not in the compressed region of mPI (Suppl. Fig 4). The absence of immune cell infiltrate is noteworthy because iron-scavenging is performed by macrophages (55, 56), and because free iron, when not adequately scavenged, can over-stimulate the innate immune response (57).

**Figure 2:**
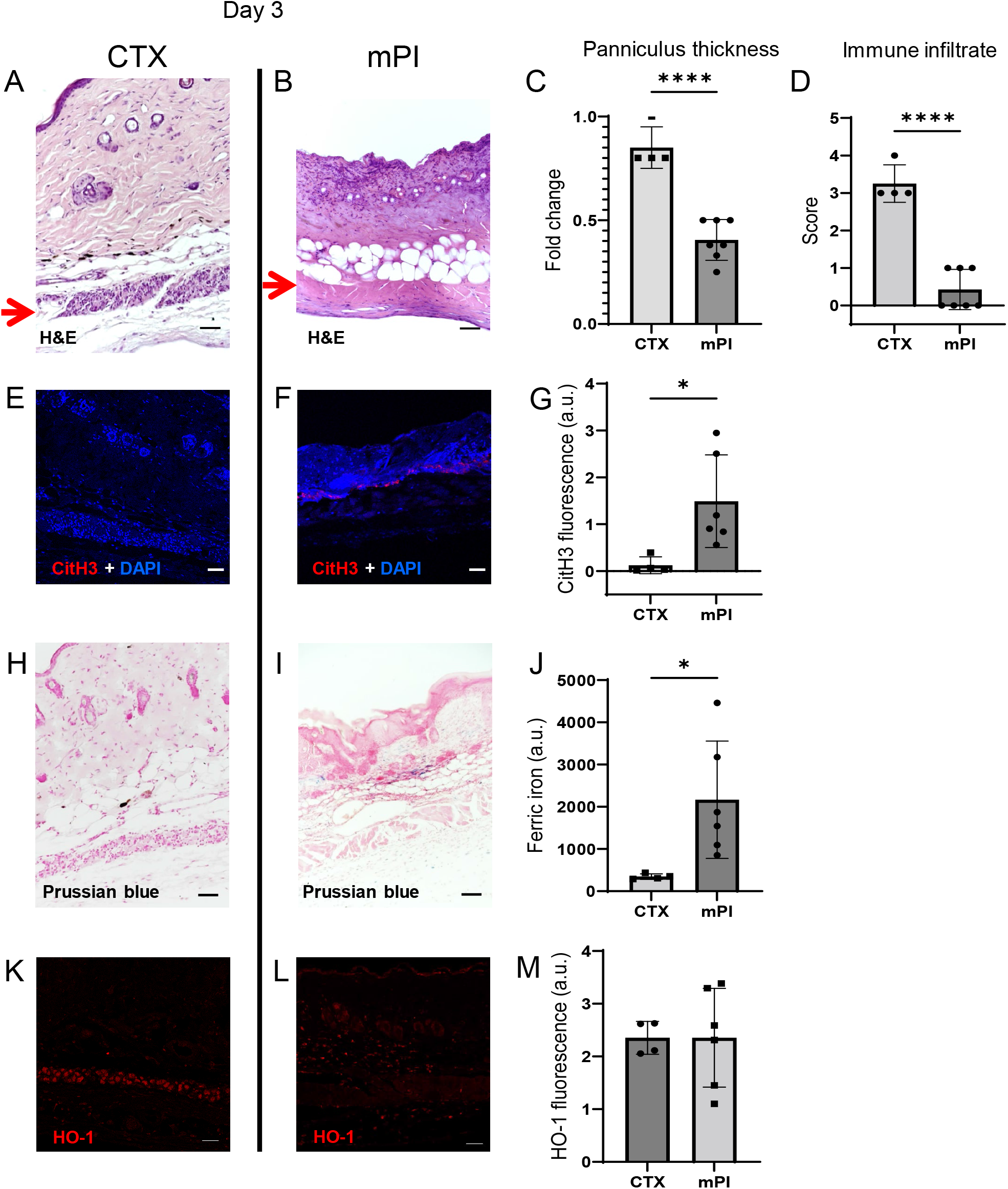
Early-stage pathologies in the injury-response of mPI (3 days after injury). (A-B) H&E-stained sections of saline-treated wound tissues, comparing magnet-induced mPI versus cardiotoxin (CTX) injury. Red arrows point to the panniculus carnosus (PC) layer. (C) Thickness of PC layer. (D) Histopathology scoring of immune infiltrate into the injured PC layer. (E-F) Immuno-staining for citrullinated histone-3 (CitH3, a marker for extracellular traps, in red) in mPI versus CTX. DNA/nuclei were co-stained blue with DAPI. (G) Quantification of CitH3 staining. (H-I) Perls’ Prussian blue iron staining (dark blue-grey), with nuclear fast red co-stain. (J) Quantification of Perls’ staining. (K-L) Immuno-staining for heme oxygenase-1 (HO-1, a marker of heme and iron) in mPI versus CTX. Note that the HO-1 positive signal in CTX injury is localized to the panniculus carnosus, and is widespread across all layers in mPI. (M) Quantification of HO-1 staining. Scale bars: 50 μm.

Another difference between mPI and CTX was in the level of citrullinated histone-3 (citH3), a marker of extracellular traps (ETs). ETs are formed when phagocytes citrullinate their histones and eject nuclear and/or mitochondrial DNA (and associated factors), which can trap and kill pathogens during infection. The process, ETosis, often kills the host cell. Extracellular traps have been observed in sickle cell ulcers (58). In day 3 wounds, levels of citrullinated histone-3 (citH3) were 10-fold higher in mPI than in CTX (Fig 2E-G; p < 0.05). Highest levels occurred near the muscle layer, such as the interface between the panniculus layer and the dWAT or dermis. This is consistent with the possibility that immune cells may have been present in the muscle layer of mPI before day 3 and then died of ETosis. Oxidative stress is a well-studied trigger of ETosis, and other stimuli include hemin and heme-activated platelets (59, 60).

### Free iron remains in wound tissues after pressure injury

Iron deposition was very high in mPI, as measured by Perls’ Prussian blue stain (Fig 2I), but iron was undetectable at the same time-point after cardiotoxin injury (Fig 2H-J; p < 0.05). Prussian blue detects accumulation of ferric Fe^3+^, typically in the form of ferritin and hemosiderin, but it only shows high levels of iron, because it is unable “to detect iron except in massive deposition” (61, 62). The blue speckles in Fig 2I are concentrated iron deposits in the extracellular matrix, and the blue ovals are iron-loaded immune cells (55, 56). Heme oxygenase-1 (HO-1) is an enzyme that performs heme degradation and serves as a marker of high heme or iron. HO-1 was expressed by mPI wound tissues, at similar levels to cardiotoxin injured tissue (Fig 2K-M; ns). However, HO-1 expression was localized to the panniculus layer after cardiotoxin injury, but it was widespread across all layers after mPI (Fig 2K-L).

### Extracellular myoglobin harms macrophage viability *in vitro*

Because immune cells were absent from the wound bed of mPI at Day 3, we asked whether free myoglobin (Mb) is cytotoxic to monocytic cells. Incubation of RAW264.7 cells in media containing 50 μg/ml myoglobin (comparable to levels in muscle lysate) caused cell death and decreased ATP production by 46% compared to cells cultured in media alone or treated with canonical M1 or M2 stimuli (Suppl. Fig 5A-B; p < 0.0001). Prior studies of erythrophagocytosis found a similar effect with hemin *in vitro* (63). However, *in vitro* studies of oxidative stress can have extensive artifacts (64, 65), so we returned to the tissue context for studying Mb-induced pathologies using a Mb-knockout mouse model.

### Myoglobin knockout mPI have less tissue death and greater immune infiltrate

To measure the contribution of myoglobin iron to mPI pathogenesis, we developed myoglobin knockout mice (Mb^−/−^) via CRISPR deletion of the entire gene from the germline. Note that prior studies of adult Mb^−/−^ mice found no obvious phenotype (66–68). Mb^−/−^ is often lethal to cardiac development during E9.5-E10.5, but some Mb^−/−^ embryos survive to term (68). Among our Mb^−/−^ that were born, all developed with normal feeding, weight gain, grooming, social behaviour, and lifespan. Deletion of Mb was confirmed by western blotting (Suppl. Fig 6A), immuno-staining (Suppl. Fig 6B-C), and DNA gel electrophoresis (Suppl. Fig 6D). With H&E staining, we detected no knockout-induced changes to the tissues of our injury model (skin, panniculus carnosus layer, or loose areolar tissue) other than increased capillary density (by 17%, p < 0.05; Suppl. Fig 6E) and increased thickness of the dWAT layer in Mb^−/−^ mice (by 43%, p < 0.05, Suppl. Fig 6E). Total iron was not significantly decreased (p = 0.066, Suppl. Fig 6F).

We compared pressure ulcers in Mb^−/−^ versus Mb^+/+^ mice (Suppl. Fig 6G) using elderly 20-month-old animals. (The mPI in elderly were similar to mPI in young, except with milder increases in pressure-induced oxidative damage, Suppl. Fig 7). At day 3 post-injury, Mb^+/+^ mice had high levels of iron (Fig 3A), which appeared in the muscle, dWAT, and dermis, including both the extracellular space and in the small numbers of infiltrating immune cells. In contrast, Mb^−/−^ mice had no detectable signal from Perls’ stain in any layer of the wound (Fig 3B-C; p < 0.001). HO-1 was also decreased by 57% in Mb^−/−^ mPI compared to Mb^+/+^ (Fig 3D-F; P < 0.05). Levels of immune cell infiltrate were 233% greater in Mb^−/−^ compared to Mb^+/+^ (Fig 3G-I; p < 0.05).

**Figure 3:**
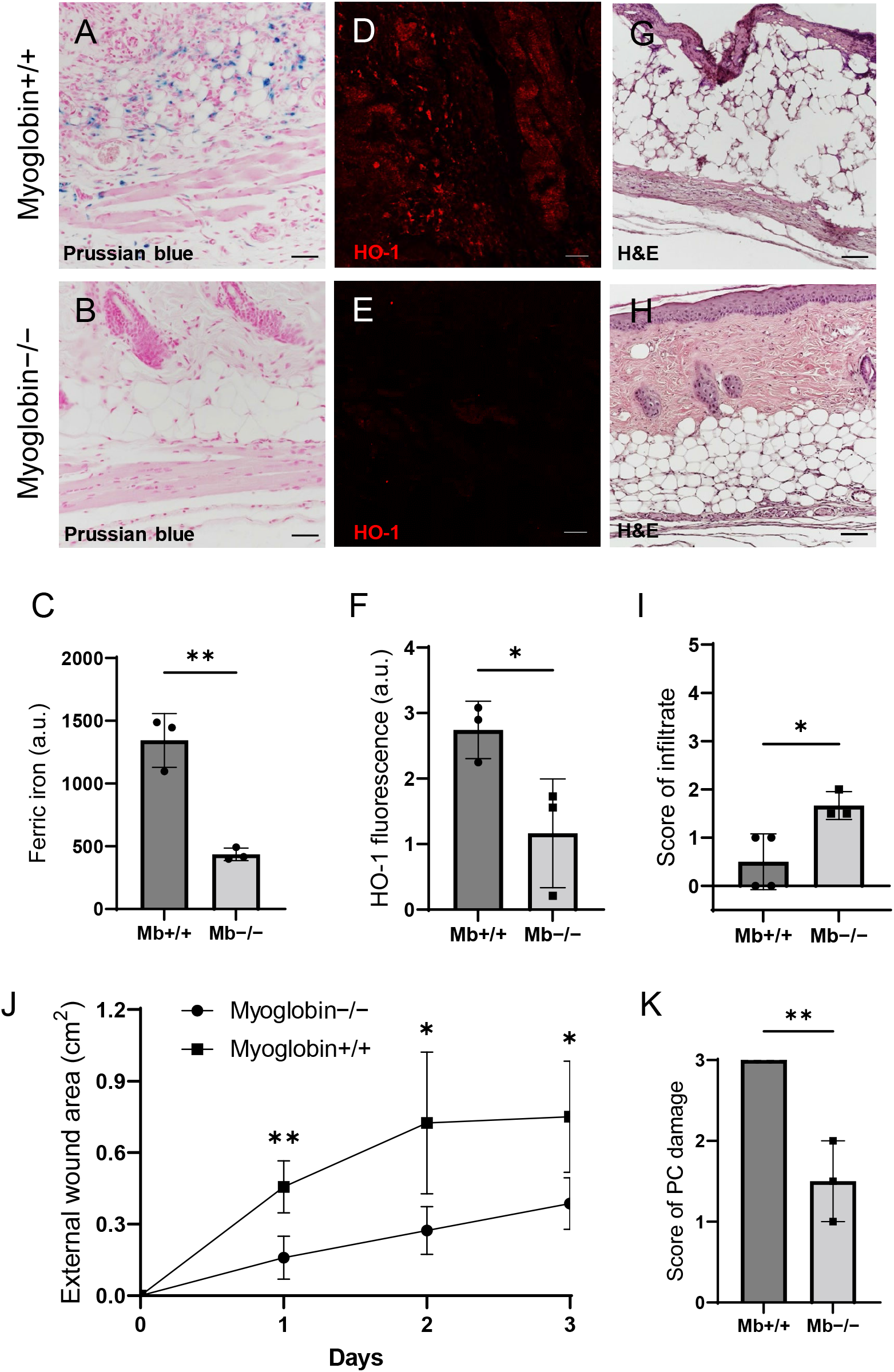
Myoglobin knockout decreased iron deposits and tissue death after mPI. (A-B) Perls’ Prussian blue iron staining. (A) Note iron deposits in the extracellular space and in immune cells of Mb+/+ wound tissue, and note that (B) Mb−/− tissues have no iron deposits in extracellular regions. (C) Quantification of Perls’ staining. (D-E) Immuno-staining for HO-1 in Mb+/+ versus Mb−/− tissues at day 3 after mPI. Note that HO-1 is elevated in all layers of Mb−/− except epidermis. (F) Quantification of HO-1 immuno-staining. (G-H) H&E-stained sections of Mb+/+ versus Mb−/− mPI. Paraffin-embedded wound sections derived from elderly Mb-wildtype mice had poor cohesiveness (compared to elderly Mb-knockout or young) and exhibited greater cracking during identical sample handling. (I) Amount of immune infiltrate, quantified by histopathology scoring on a scale of 0 to 5, performed on day 3 sections. (J) External wound area in Mb+/+ and −/− mice in the initial days following mPI from 12 mm magnets. Statistical analysis compared four wounds from two age- and sex-matched animals using a Student’s t test for each day. Consistent with these results, Suppl. Table 1 shows additional animals treated with different-sized magnets. (K) Tissue death in the PC muscle layer by histopathology scoring (3 indicates pervasive death). Statistical analyses

The wound size was smaller in Mb^−/−^ versus Mb^+/+^ (p < 0.05, measured as external area, Fig 3J and Suppl. Table 2). Histopathology scoring in the centre of the wound showed 50% decreased tissue death (p < 0.01, Fig 3G-H and Fig 3K). Oxidative damage was lower in Mb-knockout wounds: DNA damage (8-OG, 8-hydroxy-2’-deoxyguanosine) was decreased by 87% (Fig 4A-E; p < 0.05), and lipid peroxidation (measured using BODIPY 581/591) was decreased by 61% (Fig 4F-J; p < 0.05). BODIPY 581/591 is a fatty acid analogue with specific sensitivity to oxidation (69). When oxidized, its emission fluorescence shifts from 595 nm (red) to a maximal emission at 520 nm (green). The green fluorescence is what was shown in the figure. Similarly, Mb^−/−^ had roughly 56% decrease in CitH3 (Fig 4K-Q; p < 0.05). These improvements in the wound microenvironment extended beyond the muscle layer, because Mb^+/+^ wounds had high levels of BODIPY in the dWAT and dermis, and high levels of CitH3 throughout the wound. An additional measure of oxidative damage, 3-nitrotyrosine, showed the same pattern (Suppl. Fig 8A-E; 56% decrease, p < 0.05).

**Figure 4:**
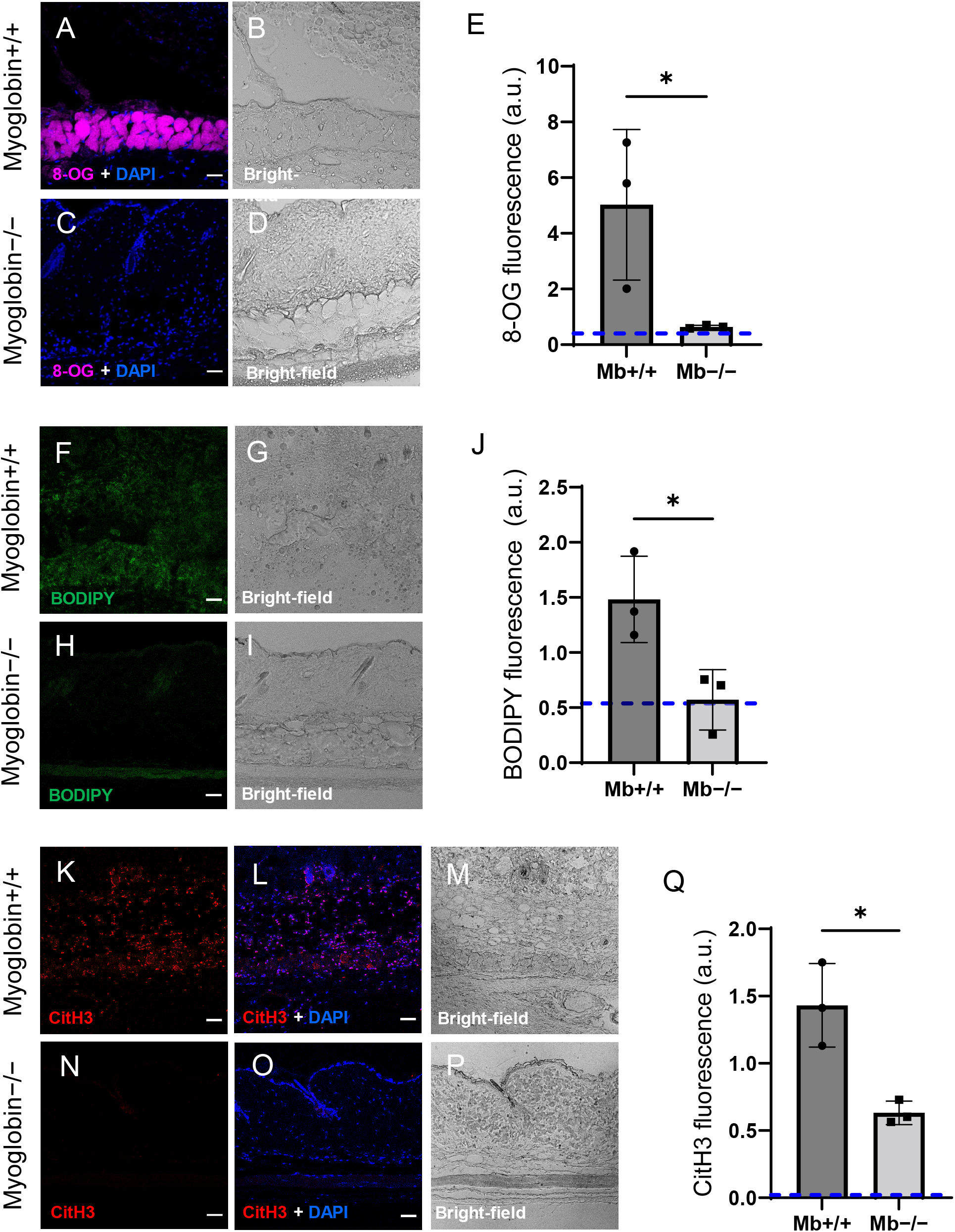
Myoglobin knockout caused a more hospitable wound environment in mPI. (A-D) Immunostaining of 8-oxaguanine (8-OG; in magenta) in Mb+/+ versus Mb−/− mPI. Nuclei were co-stained blue with DAPI. (B) and (D) are bright-field images of (A) and (C), respectively. (E) Quantification of 8-OG staining. (F-I) BODIPY staining (for lipid peroxidation) in Mb+/+ versus Mb−/−. (G) and (I) are bright-field images of (F) and (H), respectively. (J) Quantification of BODIPY staining. (K-P) Immuno-staining for CitH3 (in red) in Mb+/+ versus Mb−/−. (L, O) DNA/nuclei were co-stained blue with DAPI. (M) and (P) are bright-field images of (K) and (N), respectively. (Q) Quantification of CitH3 staining. Scale bars: 50 μm. Blue dashed lines refer to mean fluorescence intensities for uninjured dorsal skinfolds.

A panel of cytokines, chemokines and growth factors were measured in muscle homogenates of mPI from Mb^−/−^ versus Mb^+/+^ at day 3 (Suppl. Table 3). There were no significant differences except the knockout wounds had lower levels of CXCL16 (a cytokine associated with lipid peroxidation (70), Suppl. Table 3; p < 0.05) and higher levels of PAI-1 (also called Serpin E1, a protease inhibitor associated with TGFβ, Suppl. Table 3; p < 0.01). There were no significant differences in total protein between Mb^−/−^ and Mb^+/+^.

We next sought an orthogonal intervention to test the causal role of myoglobin iron. The FDA-approved iron chelation drug deferoxamine (DFO) was administered via injection under the dorsal skinfold of 5-month-old mice, starting the morning after pressure induction finished, and repeated twice daily for up 16 days (Fig 5A). DFO- or saline-treated tissues were analysed at 3, 7, 10, 40, and 90 days (Suppl. Table 4). The same cohort of saline-treated mPI were compared against saline-treated CTX in Figs 1–2 (for day 3, 10 and 40 post-injury), and compared against DFO-treated mPI in Figs 5–6.

**Figure 5:**
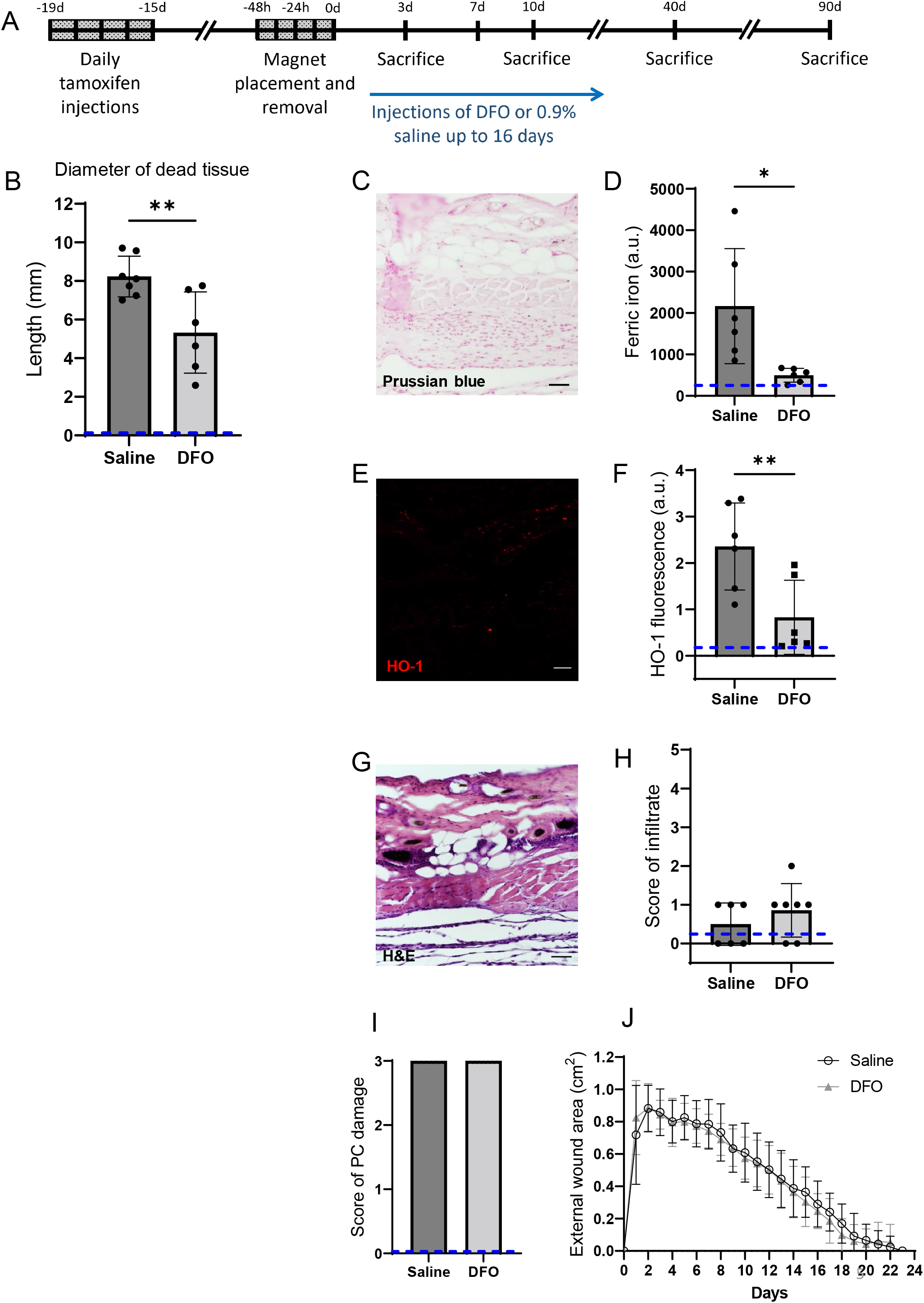
Deferoxamine (DFO) iron chelation therapy decreased iron deposits at day 3 after mPI. (A) Experimental schedule shows confetti tamoxifen-induction of fluorescence, mPI induction, treatment with DFO or saline, and tissue harvest. (B) Diameter of the dead region of the panniculus carnosus (PC) muscle in tissue sections from DFO-versus saline-treated mPI, 3 days post-injury. (C) Perls’ Prussian blue iron staining of DFO-treated wounds at the wound center. (D) Quantification of Perls’ staining, showing comparison against saline-treated mPI from Figure 2I. (E) Immunostaining of HO-1 in DFO-treated mPI. (F) Quantification of HO-1 staining, showing comparison against saline-treated mPI from Figure 2L. (G) H&E-stained sections of DFO-treated mPI at the wound center. (H) Histopathology scoring of immune infiltrate at all layers of the wound center at day 3, comparing DFO-treated versus saline-treated mPI (which appear in Fig 2B, D). (I) Confirmation that injuries were properly created, according to death of PC tissue at the center of the wound (histopathology scoring where 3 indicates pervasive tissue death). (J) Skin wound area following mPI in saline- and DFO-treated mice. Scale bars: 50 μm. Blue dashed lines refer to histology scores and mean fluorescence intensities for uninjured dorsal skinfolds. Statistical analyses were performed using a paired Student’s t test.

**Figure 6:**
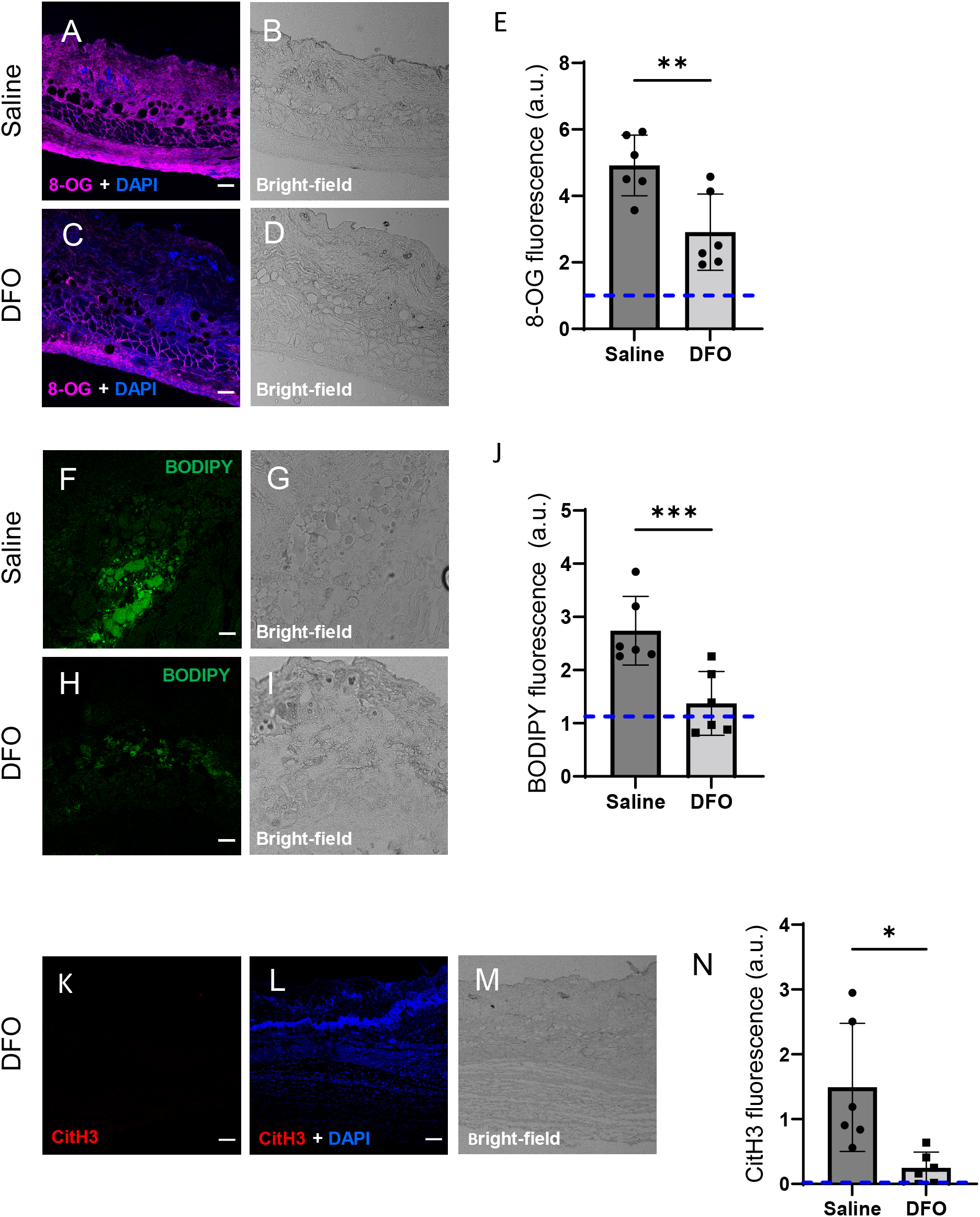
DFO treatment improved the mPI microenvironment at early time-point (day 3). (A-D) Immunostaining of 8-OG (for DNA damage) in saline-treated versus DFO-treated mPI. Nuclei were stained blue with DAPI. (B) and (D) are bright-field images of (A) and (C), respectively. (E) Quantification of 8-OG. (F-I) BODIPY staining (for lipid peroxidation) and brightfield in saline- versus DFO-treated mPI. (J) Quantification of BODIPY. (K) CitH3 immuno-staining (L) with DNA/nuclear co-stain and (M) brightfield in DFO-treated mPI at day 3. (N) Quantification of CitH3 staining in DFO-treated versus the saline-treated mPI, which was analyzed in Fig 2F-G. Scale bars: 50 μm. Blue dashed lines refer to mean fluorescence intensities for uninjured dorsal skinfolds.

### Effects of iron chelation therapy on secondary progression of the wound

Remarkably, DFO treatment caused a decrease in the amount of muscle tissue that died from the initial pressure injury, even though the pressure inductions were identical, and treatments did not begin until 12 hours after the last cycle of pressure. That is, intervention only started after deformation-injury and reperfusion-injury had already occurred. At day 3 of treatment, DFO-treated wounds had 35% smaller diameter of dead PC tissue than saline-treated (Fig 5B; p < 0.01). This observation of secondary progression confirms a prediction of our prior computational modeling (41).

### Effects of iron chelation therapy on injury response at day 3

Consistent with decreased death, DFO-treated mPI displayed 77% lower levels of iron deposition by Perls’ Prussian blue (Fig 5C-D; p < 0.05) compared to saline control (Fig 2I) at day 3. As in the elderly mPI, the young saline-treated mPI had iron accumulations in the extracellular space and immune cells of multiple layers, not just the panniculus muscle layer (Fig 2I). Similarly, levels of HO-1 were 65% decreased (Fig 5E-F and Fig 2L; p < 0.01). To confirm that wounds were properly induced, Fig 5H-I show complete tissue death at the centres of the wounds in all animals. Re-epithelialization was not affected by the subcutaneous drug (Fig 5J and Suppl. Fig 9).

DFO-treated wounds displayed less oxidative damage, as indicated by a 41% decrease in 8-OG (Fig 6A-E; p < 0.01), and 50% decrease in BODIPY 581/591 (Fig 6F-J; p < 0.001) at day 3. Change in tyrosine nitration was not significant (Suppl. Fig 10A-E). Citrullinated histone-3 (CitH3) dropped to undetectable levels in DFO-treated sections (Fig 6K-N and Fig 2F; p < 0.05). The high CitH3 found in control wounds was partially co-localized with F4/80, a pan-macrophage marker (Suppl. Fig 11A-D), but these markers may reflect debris or non-cellular localizations. Suppl. Table 5 shows additional measurements of cytokines, chemokines, and growth factors in DFO-versus saline-treated mPI at day 3 and Suppl. Table 6 shows the same analytes for Day 10. These factors showed no significant differences between DFO and saline treatments, except for CXCL16 levels, which decreased by 36% (p < 0.05) at Day 3. Other differences between saline- and DFO-treatment are shown in Suppl. Fig 11 (Suppl. Fig 11E-G).

### Effects of iron chelation therapy on immune infiltration and regeneration

At 7 days post injury, DFO-treated tissues displayed 156% greater abundance of immune infiltrate, according to brightfield and DAPI staining (Fig 7C-D, 7L; p < 0.05). Co-staining with iNOS showed strong co-localization with the infiltrate. iNOS is a marker of pro-inflammatory stimulation in many cell types, and it showed a 43% increase after DFO treatment (Fig 7A-D and Fig 7K; p < 0.05). In addition, Mer tyrosine kinase (MerTK) exhibited a 170% increase in expression in DFO-treated tissues (Suppl. Fig 12; p < 0.05). MerTK is a marker of monocytes and other specialized cell types. It has been shown to promote macrophage survival (71) and phagocytic function (specifically efferocytosis or clearance of dead cells) (72). Later at day 10, immune infiltrate was still elevated in DFO-treated wounds (p < 0.01 for histopathology and p < 0.01 for count of DAPI nuclei). Arginase-1 (Arg-1) is a marker of regenerative activity, and it promotes polyamine biosynthesis for wound healing. Arg-1 levels were two-fold higher in the wound bed at day 10 (Fig 7N; p < 0.05) and much of the day 10 infiltrate was positive for Arg-1 (Fig 7E-J and Fig 7M-O). Arg1-positive cells showed 119% greater distance of infiltration into injured tissue (Fig 7P; p < 0.01) compared to saline-treated control. Granulation was also dramatically improved. Granulation is the regenerative stage of wound healing, characterized by angiogenesis (formation of new capillaries) and proliferation of epithelial, endothelial, and fibroblast cells, along with continued presence of immune infiltrate. Angiogenesis is widely reported to increase after DFO treatment (37, 38), and DFO-treated mPI had a similar trend, evidenced by a two-fold increase in small blood vessels (Suppl. Fig 13A-G; p < 0.01).

**Figure 7:**
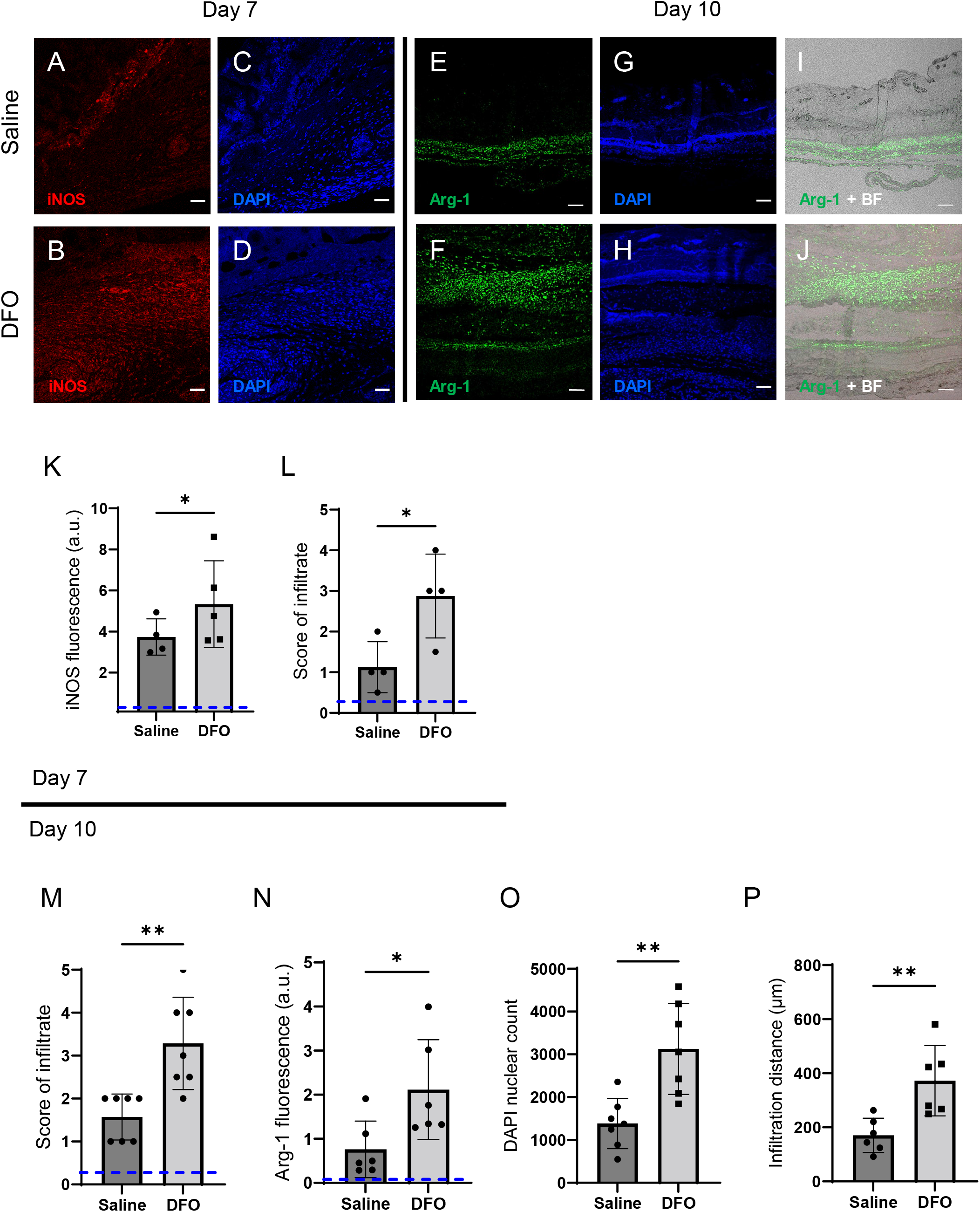
DFO treatment improved immune infiltration (7 and 10 days after mPI). (A-B) Immuno-staining of inducible nitric oxide synthase (iNOS, associated with pro-inflammatory activation) in saline-versus DFO-treated mPI at day 7 post-injury. (C-D) Nuclei are stained blue with DAPI, to show the count of infiltrating cells. (E-F) Immuno-staining of Arginase-1 in saline-versus DFO-treated mPI at day 10 post-injury. (G-H) Nuclei are co-stained blue with DAPI. (I-J) Merged brightfield and Arg-1 immuno-staining. (K) Quantification of iNOS staining at day 7 between saline- and DFO-treated. (L) Scoring of immune infiltrate into the injured tissue at day 7. (M) Scoring of immune infiltrate into the injured tissue at day 10. (N) Quantification of Arg-1 staining. (O) Count of DAPI nuclei. (P) Distance of tissue infiltrated by Arg-1+ cells in saline-versus DFO-treated tissues. Scale bars: 50 μm. Blue dashed lines refer to histology scores and mean fluorescence intensities for uninjured dorsal skinfolds.

### Quality and quantity of regeneration

DFO treatment improved the extent of muscle regeneration (Fig 8A-F and Suppl. Fig 13H). Much of this improvement was complete by day 40, when treatment improved regeneration at the wound centre (Fig 8E; p < 0.05) and wound edge (Fig 8F and Suppl. Fig 14; p < 0.05). In saline-treated wounds, myoblastic cells were observed at 40 days (Fig 8A and Fig 8C), indicating that muscle regeneration was still underway. When we extended the study to 90 days, no further increase in panniculus regeneration was detected (from 40 to 90 days) in saline-treated mPI. At the day 90 endpoint, DFO-treated wounds had significantly smaller gaps in the original region of muscle (46% smaller area of un-regenerated muscle; Fig 8G; p < 0.01) compared to saline.

**Figure 8:**
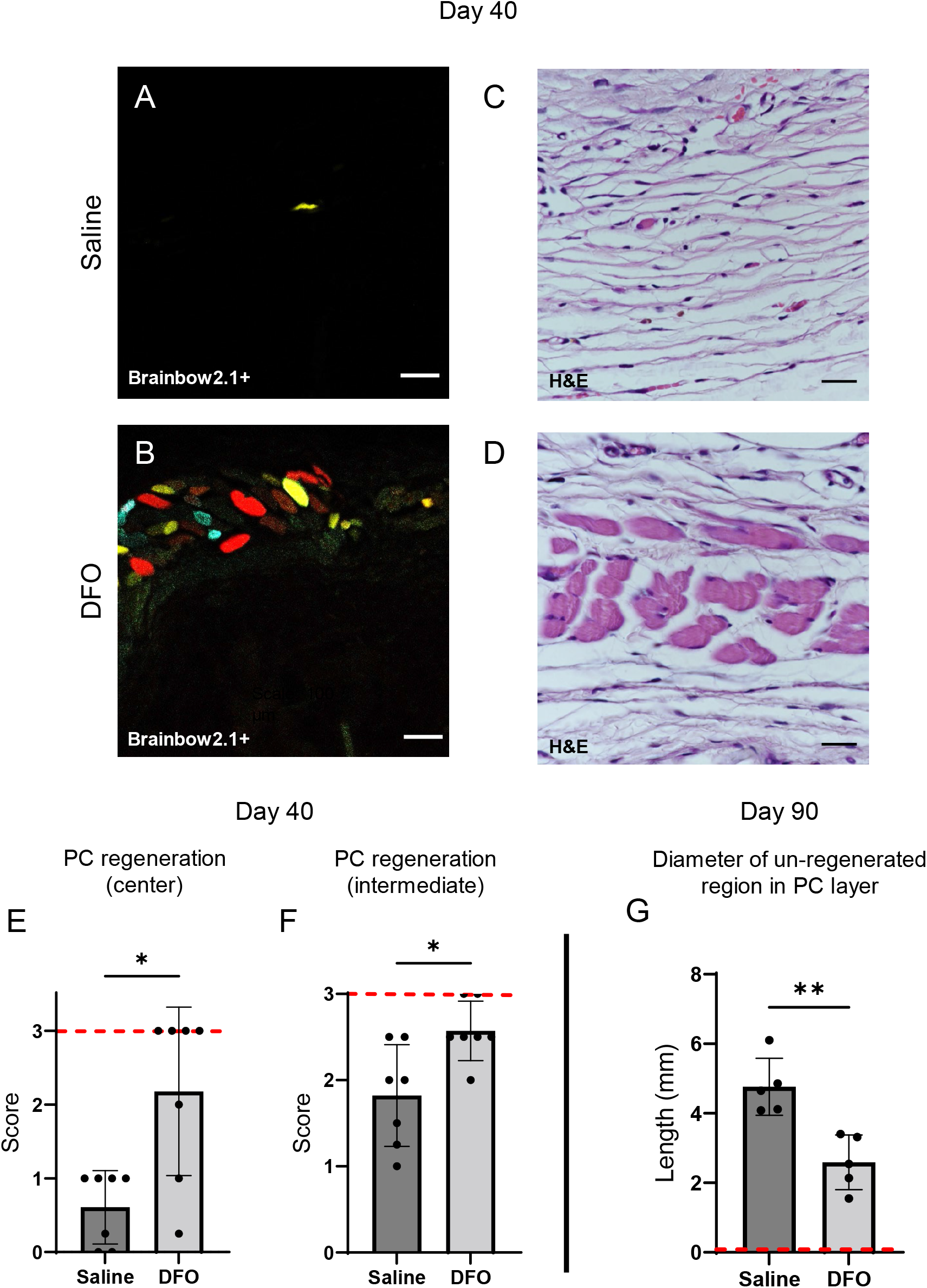
DFO increased the extent of muscle regeneration at day 40 & day 90. (A-B) Confocal fluorescent and (C-D) H&E-stained cross-sections of regenerated muscle fibers from saline- and DFO-treated mPI, 40 days post-injury. Scale bars: 100 μm. (E-F) Histological scoring of PC muscle regeneration in saline- and DFO-treated mPI versus acute CTX-injured tissues at the center and edge of the wound, 40 days post-injury. (G) Diameter of gap (unregenerated region) in PC muscle layer at day 90 post-mPI between saline- and DFO-treated wounds. Red dashed lines refer to PC regeneration scores at centre and intermediate regions after cardiotoxin injury at day 40 and diameter of unregenerated PC layer at day 90 after cardiotoxin injury.

Compared to healthy regeneration (Fig 9A-C, cardiotoxin-injured), the newly regenerated myofibers in saline-treated wounds had pathological morphology, seen from branched or split fibers, thin fibers, wavy instead of straight axes, and disjoint angles between different bundles of fibers (Fig 9D-F). Newly regenerated myofibers were distinguishable from pre-existing or never injured myofibers by their expression of fluorescent proteins, because our mice expressed the confetti transgene in Pax7-positive satellite cells and their progeny (54, 73). DFO-treated tissues displayed a far lower frequency (p < 0.001) of morphological malformations than saline-treated tissues (Fig 9G-J).

**Figure 9:**
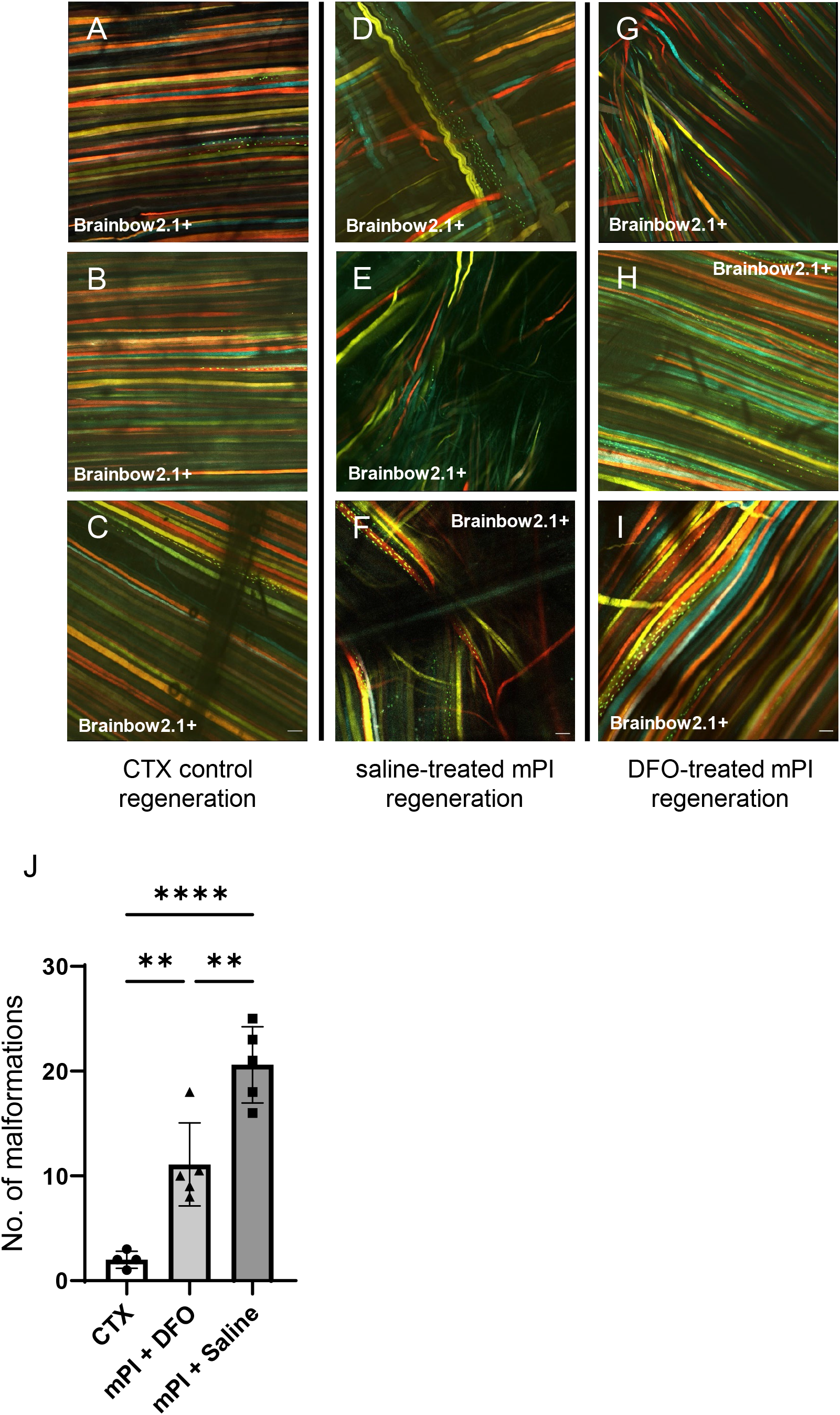
DFO improved muscle morphology. Confocal microscopy of confetti labelled muscle regeneration in ex vivo tissue blocks. Note that unlabelled tissue types such as blood vessels and hair create black shadows on top of the images. (A-C) Left column shows healthy regeneration in the control condition, toxin-induced injury, at 40 days. Experiment was halted at 40 days due to complete regeneration. (D-F) Middle column shows regenerated muscle fibers of saline-treated mPI, 90 days post-injury. Note the presence of non-parallel fibers, bent fibers, and split fibers (i.e., fibers with one or more branches). (G-I) Right column shows regenerated muscle fibers of DFO-treated mPI, 90 days post-injury. (J) Quantification of muscle fiber malformations. Scale bars: 100 μm.

## Discussion

Our study established a mouse model of muscle pressure injury (mPI) that exhibited poor healing in the absence of any infection, hyperglycemia, or old age. With blood vessels collapsed or disrupted, the mPI had dead tissue persisting in the wound bed for more than a week, and regeneration was impaired. The debris field also contained high accumulations of iron, which motivated us to study myoglobin-knockout mice and iron chelation therapy. These experiments showed that the iron was myoglobin-dependent, and the myoglobin iron caused oxidative damage. The damage was consistent with lipid peroxidation, extracellular traps, and ferroptosis. Finally, we showed that administering DFO after pressure caused decreased oxidative stress, decreased death of tissue margins, and improved muscle regeneration. The numbers of intact immune cells in mPI were positively correlated with regeneration and negatively correlated with oxidative damage, suggesting that excessive inflammation was not the primary pathology causing poor healing in these wounds.

### Induction of Mpi

used a standard magnet method in mice, but instead of studying longitudinal re-epithelialization, we used tissue section timepoints to study the thin muscle layer of panniculus carnosus, also called the cutaneous muscle or the panniculus layer. This muscle regenerated slowly and incompletely from the magnet injury (Fig 1H), even under ideal conditions of youth, health, and lack of microbial pathogens. A hole remained in the panniculus layer at 40 days (in 7 out of 7 mice) and 90 days (in 5 out of 5 mice). For example, in Figure 1D, note that the adipose layer is immediately adjacent to loose areolar tissue at the wound centre, indicating that the panniculus layer is absent. Thus, we obtained a normoglycemic non-infected animal model of impaired healing.

### Mb knockout

tested the hypothesis that myoglobin iron contributes to the pathologies of mPI. This hypothesis is based on analogy to poor-healing wounds that have hemolysis and poor drainage of hemoglobin (e.g., venous stasis), and also inspired by papers showing myoglobin is detectable in distal fluids after pressure ulcers (20, 23, 24). As expected, the iron deposits seen after wildtype mPI injury were absent from Mb^−/−^. Similarly, Mb^−/−^ had far less oxidative damage. Because Mb normally serves to supply oxygen in muscle tissue, one might expect Mb^−/−^ mice to experience increased tissue death due to the hypoxia of ischemia, but prior work showed Mb^−/−^ mice are “surprisingly well adapted” to hypoxic conditions (74), capable of withstanding adrenergic stimulation (68), exercise and oxygen flux (67). In the mPI context, carrying wildtype Mb caused roughly twice as much tissue death as Mb^−/−^. Some of that difference might be explained by developmental compensation (67) (e.g., 17% increase in capillary density), but we interpret that a large fraction of tissue death from mPI was downstream of Mb. Multiple other DAMPs including hemoglobin might have contributed to the iron accumulation and oxidative damage seen in the Mb^+/+^ case, but removing Mb was sufficient to alleviate the overload.

### The region of myglobin-related damage

extended beyond muscle. Given the potential toxicity of myoglobin during ischemia-reperfusion (66–68, 75, 76), we were not surprised that Mb knockout caused greater survival of muscle. However, we were surprised that most other layers of the wound improved too. Some types of ROS created by iron are highly reactive (e.g., hydroxyl radical from the Fenton reaction), so they act locally where they are created. Long-distance damage might arise from transport of the globin iron, or from milder ROS species, or both. Given the large effect-sizes seen in the Mb^−/−^ and DFO experiments, and given the small amount of muscle in mPI, we suspect that unappreciated mechanisms of amplification or vulnerability might be downstream of myoglobin iron in the mPI context.

### Clearance of myoglobin iron

can be seen in cardiotoxin injuries, which had destruction of muscle but no accumulation of iron, according to Perls’ stain (Fig 2J). Cardiotoxin and naniproin are snake venoms that cause myolysis (cytolysis) of myofibers (77, 78), and have been reported to spare nerve and vessel viability at the concentrations used (79, 80). One presumes that myolysis would cause myoglobin release, but continued function of the circulatory system may have allowed the myoglobin iron to be transported, diluted, and/or detoxified below the detection limit of Perls’ stain. In contrast, levels of iron in mPI were greater than the detection limit for Perls’ stain (Fig 2I), and many dysfunctions ensued. Pressure has been reported to cause disruption to arterial supply, venous clearance, and lymphatic drainage (4, 51, 52), although not always (81). Lymph vessels are occluded at very low pressure, and impaired lymphatic flow can persist after pressure is removed (82). Impaired circulation and/or impaired drainage likely caused waste factors to accumulate in the mPI, including iron. Poor drainage also occurs in chronic ulcers of venous insufficiency, lymphedema, and other vascular diseases. Toxicity from globin iron might be a shared pathology among these wounds. mPI research might also benefit analogy to known globin overload syndromes such as compartment syndrome, rhabdomyolysis, and hemoglobinopathies.

### Pressure injury prevention

usually refers to pre-injury prevention, which is to intervene with cushioning or turning before prolonged pressure occurs (39). What we studied is post-pressure prevention, because we asked whether tissue could be saved from dying, by interventions beginning 12 hours after pressure had finished. In healthy young mice, magnets were placed and removed in 12-hour cycles, as part of a two-day process of creating a pressure ulcer. (See the detailed timeline in Fig 5A.) Drug injections began on the third day after pressure began, which we refer to as Day 1 post-injury. The remarkable observation (Fig 5B) is that the amount of death on Day 3 post-injury was not constant across the tissue sections, even though the induction of the pressure ulcers was identical. mPI that received post-pressure DFO showed significantly smaller diameters of muscle death than saline-treated comparisons. Our quantification used cross-sectional tissue slides, but a 35% decreased diameter of death equates to a 58% smaller circle area of death. We conclude that there is window of opportunity to intervene and save viability of tissue, after the established mechanisms of injury have occurred (such as mechanical, hypoxic, reoxygenation, and nutrient stress), but before the full extent of secondary progression and DAMP-induced stress have propagated. In this mouse model, the opportunity occurred between 12 hr to 72 hr after off-loading.

### Secondary progression

of wounds is well-documented in thermal burns, where post-burn treatment can lessen the progression of partial-thickness burns toward full-thickness. However, secondary progression has not, to the best of our knowledge, been previously targeted for medical intervention in pressure ulcers (43). Mechanisms of secondary progression may include oxidative stress and ferroptosis (83, 84), and we cannot rule out reperfusion injury, platelet activation, extracellular traps, and DAMP-induced apoptosis/necroptosis (41).

### CitH3

a marker of extracellular traps (ETs), was elevated in the high-iron conditions (young and elderly mPI) and low in the three conditions that lacked concentrated deposits of iron (cardiotoxin, knockout, DFO-treated). This is consistent with the ability of heme, oxidative stress, or heme-activated platelets to trigger ETs (59, 60). ETs can aid in antimicrobial defense, but they are detrimental to regeneration and contribute to many disorders of sterile inflammation. Our induction of CitH3 was likely sterile as well: the mouse environment was negative for 39 categories of pathogen (Suppl. Table 7, including common dermal microbiota *P*.*aeruginosa* and *S*.*aureus*), and no bacteria could be detected in or near the wound by Gram staining.

### The persistence of dead tissue

was particularly striking in our non-infected mPI wound. Viable immune cells were essentially absent from the pressure-injured muscles of young healthy mice at Day 3. The wounds exhibited a normal spike in immune cell infiltration outside the wound margins, moving toward the wound (Suppl. Fig 4), but intact infiltrate was not seen inside the compressed region, almost as if infiltration had been halted at the boundary. Initially we questioned whether the compressed architecture of the tissue might have occluded chemotaxis, but closer inspection revealed dead fragments and markers such as CitH3 in the compressed region. Our interpretation is that immune cells (tissue-resident and/or infiltrated) may have been present but then died. We do not know whether this death was necrotic or programmed. Possible explanations for the persistence of dead tissue, in addition to poor cell survival, are decreased migration and decreased efferocytosis/phagocytosis. Diseases characterized by the presence of plaque (atherosclerosis) and ectopic tissue (endometriosis) have impaired function of phagocytes (47, 85). Other syndromes with iron overload have poor function, migration and/or survival of phagocytes (86–88). For example, in patients with thalassemia major, neutrophils and macrophages exhibit poor chemotaxis, defective lysis, and impaired phagocytosis (46, 48, 89).

### Intact immune cells

were present at Day 7 and Day 10 of mPI. DFO increased their abundance, especially in the loose areolar tissue, but also in muscle, fat, and skin (Fig 7L-M). One theory of non-healing wounds is that excessive inflammation causes oxidative damage and blocks progress of granulation. Our mPI observations are not well aligned with that theory because DFO caused an increase in the immune infiltrate and iNOS staining, but it ***decreased*** the oxidative damage and the size of the hole. Another theory is that non-healing wounds lack pro-regenerative M2 macrophages. Nothing observed about mPI in this first study would contradict that theory. Our work is also consistent with prior work that found iron-scavenging macrophages from an iron-loaded tissue displayed both inflammatory (iNOS, IFNγ) and alternately-activated (Arg1, IL10) markers (90). Prior studies also found that environmental iron can cause M2 macrophages to decrease (91) or increase (92), depending on dose/context. Future studies can use our mPI timeline to design an immuno-biology study that isolates mPI immune cells for detailed analysis.

### Surface visibility of dead tissue

was delayed until competent inflammation and granulation pushed it upward. This delay might give a false impression that slough was created by infection or inflammation. The slough detached spontaneously by Day 15, but as it was pushed up, the slough remained attached to the healthy margins and bent the margins upward (Fig 1E and Suppl. Fig 13). Since healthy margins organize the geometry of regeneration, distorting the margins might impair regeneration. Non-healing wounds are sometimes believed to have excess proteolysis, but mPI raise the question of whether increased proteolytic activity might help prevent the extrusion process from distorting the healthy margins.

### The toxicity of slough

is a topic of clinical debate. In our mPI, re-epithelialization proceeded beneath the necro-slough and proliferative granulation tissue was adjacent to its centre, so we conclude it was not cytotoxic. This is consistent with clinical practice to avoid debridement of an intact eschar unless there are signs of infection (93). Interestingly, our slough arose from a mass that was probably toxic when it formed. Toxicity at early timepoints is inferred from the oxidative damage at day 3, and indirectly from the phenomenon of secondary progression. It is possible that toxicity could disappear spontaneously if ROS are driven by an energy source that gets depleted (e.g., hydrogen peroxide, ATP, mitochondrial membrane potentials).

### Muscle regeneration

improved with DFO treatment, evidenced by smaller holes in the muscle layer at multiple timepoints, and improved muscle morphology at the endpoint. The limited muscle regeneration that did occur in mPI without DFO displayed frequent myofiber defects such as branching and non-parallel alignment. Split fibers are considered defective regeneration (94) and are especially vulnerable to re-injury (95). Loss of the panniculus muscle layer causes the skin and adipose to experience harsher stress and strain (54, 96), even though the panniculus is a thin component of the tissue. Previous studies of muscle pressure injury considered larger muscles (97–101), and the impact of DFO should be retesting in other geometries. The panniculus layer is highly relevant to human pressure injuries because humans have a panniculus layer at the heel (54, 102), and the heel is one of the most common locations for high-stage pressure injuries.

### Skin regeneration

was not significantly affected by subcutaneous DFO (Fig 5J and Suppl. Fig 9). Previous work found DFO administered via a transdermal patch accelerated skin closure (37), but these studies used topical delivery, as well as using a different mouse background. Other work studied the antioxidant and angiogenic benefits of DFO in preclinical models of ischemia-reperfusion injury (103–106) and related conditions (107). We observe similar effects in muscle.

### Caveats

to this work include the following. Mouse wounds differ from human in several obvious ways, such as having denser hair follicle coverage, faster kinetics of DFO, different timing and ratio of phagocytic cells (108), and higher propensity for wound contraction. Although mice can heal skin wounds by contraction, our multi-layer mPI exhibited minimal contraction, as seen from prior work (54) and from the hole in the panniculus muscle layer at day 90. To decrease the impact of hair follicles on results, we placed magnets on regions of skin that were in the resting phase of the hair cycle. Another limitation of our study is the limited number of myoglobin knockout mice (n = 3) compared with n = 7 for key timepoints of Mb^+/+^ analysis. This was necessary because Mb germline knockout had high rates of embryonic lethality. Despite the smaller sample size, several readouts were statistically significant.

## Conclusion

Myoglobin iron contributed to the size, severity, oxidative damage, and poor healing of mPI. Deferoxamine injections prevented death of tissue at the margins of the wound, when administered starting 12h after pressure was removed. Intervention to block secondary progression is a dramatic new opportunity for pressure injuries, like burns or strokes, to be stabilized shortly after injury, to decrease further loss of salvageable tissue. Iron chelation therapy has side-effects when used systemically, so muscle wounds might be safer than vascular wounds for future research on human biodistribution (in conjunction with dressings and polymer scaffolds). Also, deferoxamine led to higher quality of muscle, with straighter and more parallel muscle fibers, which should improve muscle viability. Healthy muscle is a crucial defence against wound recurrence, when a patient eventually puts weight on the same location again.

## Methods

### Mice

Animal experiments were approved by the institutional animal care and use committee (IACUC SHS/2016/1257) of SingHealth, Singapore. To conditionally label Pax7+ muscle satellite stem cells in C57BL6 mice, the Pax7-Cre-ER^T2^ mouse (JAX stock #012476, Jackson Laboratory, ME, USA) was crossed with the Brainbow2.1 (confetti; JAX stock #017492, Jackson Laboratory, ME, USA) mouse. Pax7-Cre-ERT2 provides the Cre-ERT2 transgene downstream of the Pax7 stop codon, thereby limiting the Cre-ERT2 expression to Pax7+ cells. The Cre-ER^T2^ system provides tamoxifen-dependent induction of Cre recombinase, so that affected cells carry heritable rather than transient modification. Upon tamoxifen treatment, the Cre induction causes recombination of the confetti construct at its loxP loci, leading to gene expression of one of the four fluorescent proteins in the construct. The fluorescent proteins are mCerulean (CFP^mem^), hrGFP II (GFP^nuc^), mYFP (YFP^cyt^) and tdimer2(12) (RFP^cyt^). CFP^mem^ contains a localization sequence enabling its transport to the myofibre membrane (sarcolemma) while GFP^nuc^ contains a nuclear localization sequence. YFP^cyt^ and RFP^cyt^ have no localization sequences and are expected to localize to the cytoplasm. Experimental mice include 5-month-old adult mice and 20-month-old elderly mice.

Myoglobin knockout mice (homozygous in the germline) were created by Cyagen (CA, US) using CRISPR/Cas9 nuclease-mediated genome editing (109, 110). 20-month-old mice (n = 3) were used in the knockout experiments. The myoglobin knockout mice were not injected with tamoxifen.

Mice were euthanized via CO_2_ inhalation, followed by cervical dislocation, at timepoints of three, seven, ten, forty, and ninety days following completion of the injury protocol (pressure or toxin). Pressure injury via magnetic compression created two wounds (right and left) on the dorsum of each mouse. Similarly, two cardiotoxin injections were performed on each mouse at the right and left dorsal skinfold. Tissue samples were isolated, and one wound tissue (from either pressure or toxin) was fixed in 4% paraformaldehyde at 4°C for eight hours, and then embedded in paraffin. The remaining wound tissue was snap-frozen with isopentane in a liquid nitrogen bath.

### Murine pressure injury model

Mice were shaved and remaining hair was removed by hair removal cream (Veet, Reckitt, Slough, England) prior to injury. Muscle pressure ulcers were created in 4 to 5-month-old transgenic mice by applying a pair of ceramic magnets (Magnetic Source, Castle Rock, CO, part number: CD14C, grade 8) to the dorsal skinfold. The magnets were 5 mm thick and 12 mm in diameter, with an average weight of 2.7 g and pulling force of 640 g. One of the elderly mice lacked sufficient skinfold for 12 mm magnets, so in that mouse and in its age- and sex-matched control, we used 5 mm magnets (neodymium magnets with a 1 mm spacer on one side; Liftontech Supreme Pte Ltd, Singapore). Because hair follicle (HF) stem cells contribute to wound healing, we attempted to minimize the HF differences by synchronizing the hair growth cycle and applying injury only to skin with HFs in telogen phase (non-pigmented skin). In the case where pigmented skin could not be avoided during magnet placement, i.e., one of the two wounds fell on skin with active hair cycle, that half was excluded from further analysis. Pressure ulcer induction was performed in two cycles. Each cycle was made up of a 12-hour period of magnet placement followed by a 12-hour period without magnets (Suppl. Fig 1A). This procedure induces two pressure wounds on the back of the mouse, on the left and right side of the dorsal skinfold. The dorsal skinfold includes skin, adipose tissue, panniculus carnosus muscle, and loose areolar tissue. The mice were given an analgesic (buprenorphine at 0.1 mg/kg; Bupredyne, Jurox Animal Health, New Zealand) prior to magnet placement and again prior to magnet removal. Time points post-injury are measured from the end of last magnet cycle. No dressings were used and no debridements were performed.

To treat the pressure injuries, mice were subcutaneously injected with deferoxamine (DFO) while control mice were injected with 0.9% saline (n = 7 each for day 3, 10 and 40 time-points, n = 4 for day 7 and n = 5 for day 90; Suppl. Table 3). The study was initially conducted using 3, 10 and 40 days. When the day 7 and day 90 timepoints were added to the study, we already knew that DFO treatment would give a large effect-size. The updated effect-size caused our power calculation to give a smaller sample-size for day 7 and day 90. The day 7 timepoint was added later because of the dramatic changes observed between day 3 and day 10, and the day 90 timepoint was added later because day 40 mPI showed signs of ongoing regeneration (myoblastic cells and immature myofibers), indicating that steady-state had not yet been reached at day 40.

DFO was administered twice per day at 30mg/kg body weight (or 0.9% saline for control animals) for 16 days, or until mouse sacrifice, whichever was sooner. Under aesthesia, the dorsal skinfold (cranial to the wound) was pulled away from the spine to make a “tent,” and the needle was inserted into the tent (over the spine, near the left and right wounds). The first DFO treatment was given 12 hours after completion of the second and final cycle of pressure (Fig 5A). The dosing rationale was as follows: the recommended dose of DFO for iron overload in human patients is 20-60 mg/kg per day (maximum 80 mg/kg; (111)). Mice metabolize DFO faster than humans, so we took the 60 mg/kg per day human dosing and divided it into two half-doses per day.

### Murine cardiotoxin injury model

To induce an acute injury in the panniculus carnosus muscle in the skinfold, mice were shaved and fur removed by hair removal cream (Veet, Reckitt, Slough, England) prior to injury. The mice were anaesthetized and 30 μl of toxin at 10 μM concentration was injected intramuscularly into the panniculus carnosus of the dorsal skinfold of each wound (left and right). Because commercial distribution of cardiotoxin from *Naja mossambica* was discontinued, some mice were injected with naniproin instead, while the term “cardiotoxin” has been used for the entire cohort. Naniproin is a *Naja nigricollis* homologue of cardiotoxin with 95% sequence identity. Naniproin induces myolysis comparable to cardiotoxin. All toxin-injected mice were given an analgesic (buprenorphine at 0.1 mg/kg; Bupredyne, Jurox Animal Health, New Zealand) prior to injection. To control for the post-injury injections given to mPI mice, the toxin-injured mice were injected subcutaneously with 0.9% saline (n = 7) twice daily for 16 days or until mouse sacrifice, whichever was sooner (Suppl. Fig 1B). Tissues were harvested at day 3, 10 or 40 (n = 4). Uninjured healthy tissues were also collected as controls (n = 4).

### Histopathology scoring of H&E-stained sections

H&E-stained slides were blinded and scored on the extent of tissue death, immune infiltration, granulation, and regeneration. The scoring for tissue death was defined as follows: 0—healthy tissue with zero or minimal death (<10% tissue area dead); 1—mild death (11-33%); 2—moderate death (34-67%); 3—extensive tissue death (>67%). Death was identified by karyolysis, karyorrhexis, and acidification (eosinification). Scoring for tissue regeneration was defined as follows: 0 (minimal i.e., <10% regenerated), 1 (mild i.e., 11-33% regenerated), 2 (moderate i.e., 34-67% regenerated) to 3 (extensive i.e., >67% regenerated). Scoring for granulation was defined as follows: 0—normal tissue with neither granulation nor neo-angiogenesis; 1—minimal granulation (less than 10% of tissue area); 2—mild granulation (10% to 25% of tissue area); 3—moderate granulation (26% to 50% of tissue area); 4—high granulation (51% to 75% of tissue area); and 5—extensive granulation (more than 75% of tissue area consisting of granulation tissue). Likewise, the scores for immune infiltration were defined as follows: 0—normal tissue without immune infiltration; 1—minimal immune infiltration (in less than 10% of tissue); 2— mild immune infiltration (in 10% to 25% of tissue); 3— moderate immune infiltration (in 26% to 50% of tissue); 4—high immune infiltration (51% to 75% of tissue area); and 5—extensive immune infiltration (more than 75% of tissue infiltrated by immune cells). Therefore, on the scales for tissue death, regeneration, granulation and immune infiltration, an uninjured tissue would receive scores of 0, 3, 0, and 0, respectively. Scoring was performed for all treatments and all time-points. The scoring of death, immune infiltration and regeneration was performed for the overall wound and each tissue layer.

### Immunofluorescence staining

10 μm fixed paraffin-embedded sections or cryosections were blocked with 10% normal serum and permeabilized with 0.2% Tween 20. Staining was performed using antibodies against 8-Oxoguanine (8-OG or 8-hydroxy-2’-deoxyguanosine; Abcam, Cambridge, UK, ab206461), F4/80 (Santa Cruz Biotechnology Inc, TX, USA, sc-52664) Arginase-1 (Arg1; Abcam, ab60176 and Proteintech Group Inc, IL, USA, 16001-1-AP), MerTK (Abcam, ab95925), inducible nitric oxide synthase (iNOS; Cell Signalling Technology, MA, USA, #13120), citrullinated histone H3 (citH3; Abcam, ab5103), nitrotyrosine (Abcam, ab7048), and myoglobin (Cell Signalling Technology, 25919S). For detection, we used Alexa-Fluor 488-, 594- and 647-conjugated secondary antibodies (Abcam, ab150129, ab150064, ab150075 respectively) raised in appropriate species for the experiments. The slides were mounted with Vectashield Hardset with DAPI (Vector Laboratories, CA, USA) and images were acquired on a Leica TCS SP8 confocal microscope (Leica Microsystems, Wetzlar, Germany) and analysed using LAS X software. For quantification, each wound consists of 512 × 512 image frames, covering all layers of the wound, and the number of frames depends on the size of the wound. Each 512 × 512 image frame was thresholded using Fiji (ImageJ) as outlined by Shihan et al. (112), and the mean fluorescence intensity was computed. The mean intensity for each wound is computed by taking the mean of the frames. Text description has been added if specific layers were disproportionately responsible for the intensity.

### Statistical analyses

In the analysis of DFO treatment in adult 5-month-old mice at multiple timepoints, treated mice were paired with age-and-sex-matched controls. Therefore, significance was measured using a paired test (two-tailed Student’s t-test for mPI+DFO versus mPI+saline). Mice were not paired for the other comparisons (i.e., adult 5-month-old mPI+saline versus CTX+saline cohorts2, and elderly 20-month-old Mb^−/−^ mPI+saline versus Mb^+/+^ mPI+saline cohorts), and statistical significance was analyzed using an unpaired two-tailed Student’s t-test. For multiple comparisons in Figure 9, in a one-way ANOVA (analysis of variance) was followed by the Tukey post-hoc test. For multiple comparisons in the Luminex assays (Supplementary Tables 2, 4, 5), the Student’s t-test was performed for each analyte, followed by the Bonferroni-Dunn correction for multiple hypothesis testing. Tests and plots were generated by GraphPad Prism (version 9.0.0 for Windows, GraphPad Software, CA, USA). An asterisk (*) refers to a p value less than 0.05, (**) means p < 0.01, (***) means p < 0.001 and (****) means p < 0.0001. “ns” means not significant.

## Data availability

Primary images, data and quantification spreadsheets and mice numbers have been uploaded to Zenodo at doi: 10.5281/zenodo.7069780.

## Author contributions

NJMN: Investigation, Data analysis, Writing. HH: Methodology, Investigation. JJ: Investigation, Project administration. NHH: Sample processing. RB: Methodology. LTK: Conceptualization, Funding acquisition, Supervision, Writing.

## Acknowledgements

We acknowledge funding from the Singapore Ministry of Health’s National Medical Research Council (NMRC/OFIRG/0007/2016), and also from the Singapore Ministry of Education’s Tier2 Grant (MOE2019-T2-1-138) and the National Research Foundation, Prime Minister’s Office, Singapore under its Campus for Research Excellence and Technological Enterprise (CREATE) programme, through Singapore MIT Alliance for Research and Technology (SMART): Critical Analytics for Manufacturing Personalised-Medicine (CAMP) Inter-Disciplinary Research Group.

We thank Manjunatha Kini and Koh Cho Yeow for naniproin; Ann-Marie Chacko, Tham Jing Yang and Sarina Heng for detection methods; N. Suhas Jagannathan for image analysis; Aadya S. Deshpande, Khadijah Zulkifli, Colin Nicholas Sng and Korn Laongkul for assistance; Peter T. C. So, Paul Matsudaira, Jasmine Chin, Mynn Varela, Ruklanthi de Alwis, the Mechanobiology Institute (Singapore) Microscopy facility, and the SingHealth Advanced BioImaging facility for advice.

## Notes

The authors have declared that no conflict of interest exists.

https://10.5281/zenodo.7069780

### Competing Interest Statement

The authors have declared no competing interest.

### Summary of Updates

Text was amended so that all instances of 'failure of phagocytosis' will be written as 'persistence of dead tissue.' The title is changed to omit the term 'phagocytic dysfunction.' The discussion section has been shortened by 33%. Figure 2L and Figure 5E were revised. Suppl. Table 1 and Suppl. Fig 2 were added to show surface area and vascular comparisons between injuries from CTX and mPI. Supplemental files were updated. Acknowledgements and References have been updated.

## References

1. Girouard K, Harrison MB, VanDenKerkof E. The symptom of pain with pressure ulcers: a review of the literature. Ostomy Wound Manage. 2008;54(6):8.

2. Padula WV, Delarmente BA. The national cost of hospital-acquired pressure injuries in the United States. Int Wound J. 2019;16(3):634–40.

3. Wassel CL, Delhougne G, Gayle JA, Dreyfus J, Larson B. Risk of readmissions, mortality, and hospital-acquired conditions across hospital-acquired pressure injury (HAPI) stages in a US National Hospital Discharge database. Int Wound J. 2020;17(6):1924–34.

4. Gray RJ, Voegeli D, Bader DL. Features of lymphatic dysfunction in compressed skin tissues - Implications in pressure ulcer aetiology. J Tissue Viability. 2016;25(1):26–31.

5. Mervis JS, Phillips TJ. Pressure ulcers: Prevention and management. J Am Acad Dermatol. 2019;81(4):893–902.

6. Bouten CV, Oomens CW, Baaijens FP, Bader DL. The etiology of pressure ulcers: skin deep or muscle bound? Arch Phys Med Rehabil. 2003;84(4):616–9.

7. Preston A, Rao A, Strauss R, Stamm R, Zalman D. Deep tissue pressure injury: A clinical review. Am J Nurs. 2017;117(5):50–7.

8. Goldman DW, Breyer III RJ, Yeh D, Brockner-Ryan BA, Alayash AI. Acellular hemoglobin-mediated oxidative stress toward endothelium: a role for ferryl iron. Am J Physiol. 1998;275(3):H1046–53.

9. Reeder BJ, Wilson MT. Hemoglobin and myoglobin associated oxidative stress: from molecular mechanisms to disease states. Curr Med Chem. 2005;12(23):2741–51.

10. Kato GJ, Steinberg MH, Gladwin MT. Intravascular hemolysis and the pathophysiology of sickle cell disease. J Clin Invest. 2017;127(3):750–60.

11. Nader E, Romana M, Connes P. The Red Blood Cell-Inflammation Vicious Circle in Sickle Cell Disease. Front Immunol. 2020;11(454).

12. Mendonça R, Silveira AAA, Conran N. Red cell DAMPs and inflammation. Inflamm Res. 2016;65(9):665–78.

13. Bozza MT, Jeney V. Pro-inflammatory actions of heme and other hemoglobin-derived DAMPs. Front Immunol. 2020;11:1323.

14. Cherayil BJ. The role of iron in the immune response to bacterial infection. Immunol Res. 2011;50(1):1–9.

15. Bogdan C. Oxidative burst without phagocytes: the role of respiratory proteins. Nat Immunol. 2007;8:1029–31.

16. Jiang N, Tan NS, Ho B, Ding JL. Respiratory protein–generated reactive oxygen species as an antimicrobial strategy. Nat Immunol. 2007;8:1114–22.

17. Bahl N, Winarsih I, Tucker-Kellogg L, Ding JL. Extracellular haemoglobin upregulates and binds to tissue factor on macrophages: implications for coagulation and oxidative stress. Thromb Haemost. 2014;111(1):67–78.

18. Bahl N, Du R, Winarsih I, Ho B, Tucker-Kellogg L, Tidor B, et al. Delineation of lipopolysaccharide (LPS)-binding sites on hemoglobin: from in silico predictions to biophysical characterization. J Biol Chem. 2011;286(43):37793–803.

19. Tchanque-Fossuo CN, Dahle SE, Buchman SR, Isseroff RR. Deferoxamine: potential novel topical therapeutic for chronic wounds. Br J Dermatol. 2017;176(4):1056–9.

20. Traa WA, Strijkers GJ, Bader DL, Oomens CWJ. Myoglobin and troponin concentrations are increased in early stage deep tissue injury. J Mech Behav Biomed Mater. 2019;92:50–7.

21. Makhsous M, Lin F, Pandya A, Pandya MS, Chadwick CC. Elevation in the serum and urine concentration of injury-related molecules after the formation of deep tissue injury in a rat spinal cord injury pressure ulcer model. PM R. 2010;2(11):1063–5.

22. Loerakker S, Huisman ES, Seelen HAM, Glatz JFC, Baaijens FPT, Oomens CWJ, et al. Plasma variations of biomarkers for muscle damage in male nondisabled and spinal cord injured subjects. J Rehabil Res Dev. 2012;49(3):361–72.

23. Levine JM. Rhabdomyolysis in association with acute pressure sore. J Am Geriatr Soc. 1993;41(8):870–2.

24. Traa WA. Evaluating aetiological processes for monitoring and detection of deep tissue injury: Technische Universiteit Eindhoven; 2019.

25. Reeder BJ, Hider RC, Wilson MT. Iron chelators can protect against oxidative stress through ferryl heme reduction. Free Radic Biol Med. 2008;44(3):264–73.

26. Rojkind M, Domínguez-Rosales JA, Nieto N, Greenwel P. Role of hydrogen peroxide and oxidative stress in healing responses. Cell Mol Life Sci. 2002;59(11):1872–91.

27. Kapralov A, Vlasova II, Feng W, Maeda A, Walson K, Tyurin VA, et al. Peroxidase activity of hemoglobin-haptoglobin complexes: covalent aggregation and oxidative stress in plasma and macrophages. J Biol Chem. 2009;284(44):30395–407.

28. Plotnikov EY, Chupyrkina AA, Pevzner IB, Isaev NK, Zorov DB. Myoglobin causes oxidative stress, increase of NO production and dysfunction of kidney’s mitochondria. Biochim Biophys Acta. 2009;1792(8):796–803.

29. Osawa Y, Williams MS. Covalent crosslinking of the heme prosthetic group to myoglobin by H2O2: toxicological implications. Free Radical Biology and Medicine. 1996;21:35–41.

30. Boutaud O, Roberts LJnd. Mechanism-based therapeutic approaches to rhabdomyolysis-induced renal failure. Free Radic Biol Med. 2011;51(5):1062–7.

31. Dixon SJ, Lemberg KM, Lamprecht MR, Skouta R, Zaitsev EM, Gleason CE, et al. Ferroptosis: an iron-dependent form of nonapoptotic cell death. Cell. 2012 149(5):1060–72.

32. Velasquez J, Wray AA. Deferoxamine. Treasure Island (FL): StatPearls Publishing; 2021 May 31.

33. Karnon J, Tolley K, Vieira J, Chandiwana D. Lifetime cost-utility analyses of deferasirox in beta-thalassaemia patients with chronic iron overload: a UK perspective. Clin Drug Investig. 2012;32(12):805–15.

34. Morel I, Cillard J, Lescoat G, Sergent O, Pasdeloup N, Ocaktan AZ, et al. Antioxidant and free radical scavenging activities of the iron chelators pyoverdin and hydroxypyrid-4-ones in iron-loaded hepatocyte cultures: comparison of their mechanism of protection with that of desferrioxamine. Free Radic Biol Med. 1992;13(5):499–508.

35. Vanek T, Kohli A. Biochemistry, Myoglobin. Treasure Island (FL): StatPearls Publishing; 2021.

36. Xiao H, Gu Z, Wang G, Zhao T. The possible mechanisms underlying the impairment of HIF-1α pathway signaling in hyperglycemia and the beneficial effects of certain therapies. Int J Med Sci. 2013;10(10):1412–21.

37. Duscher D, Neofytou E, Wong VW, Maan ZN, Rennert RC, Inayathullah M, et al. Transdermal deferoxamine prevents pressure-induced diabetic ulcers. PNAS. 2015;112(1):94–9.

38. Holden P, Nair LS. Deferoxamine: An angiogenic and antioxidant molecule for tissue regeneration. Tissue Eng Part B Rev. 2019;25(6):461–70.

39. Sundin BM, Hussein MA, Glasofer S, El-Falaky MH, Abdel-Aleem SM, Sachse RE, et al. The role of allopurinol and deferoxamine in preventing pressure ulcers in pigs. Plast Reconstr Surg. 2000;105(4):1408–21.

40. Soloniuk DS, Perkins E, Wilson JR. Use of allopurinol and deferoxamine in cellular protection during ischemia. Surg Neurol. 1992;38(2):110–3.

41. Jagannathan NS, Tucker-Kellogg L. Membrane permeability during pressure ulcer formation: A computational model of dynamic competition between cytoskeletal damage and repair. J Biomech. 2016;49(8):1311–20.

42. Stadler I, Zhang RY, Oskoui P, Whittaker MB, Lanzafame RJ. Development of a simple, noninvasive, clinically relevant model of pressure ulcers in the mouse. Journal of Investigative Surgery. 2004;17(4):221–7.

43. European Pressure Ulcer Advisory Panel, National Pressure Injury Advisory Panel, Pan Pacific Pressure Injury Alliance. Prevention and treatment of pressure ulcers/injuries: Clinical practice guideline. The international guideline. Haesler E, editor: EPUAP/NPIAP/PPPIA; 2019.

44. Ferris AE, Harding KG. An overview of the relationship between anaemia, iron, and venous leg ulcers. Int Wound J. 2019;16(6):1323–9.

45. Sindrilaru A, Peters T, Wieschalka S, Baican C, Baican A, Peter H, et al. An unrestrained proinflammatory M1 macrophage population induced by iron impairs wound healing in humans and mice. J Clin Invest. 2011;121(3):985–97.

46. Ballart IJ, Estevez ME, Sen L, Diez RA, Giuntoli J, de Miani SA, et al. Progressive dysfunction of monocytes associated with iron overload and age in patients with thalassemia major. Blood. 1986;67(1):105–9.

47. Liu YY, Liu YK, Hu WT, Tang LL, Sheng YR, Wei CY, et al. Elevated heme impairs macrophage phagocytosis in endometriosis. Reproduction. 2019;158(3):257–66.

48. Martins R, Maier J, Gorki AD, Huber KVM, Sharif O, Starkl P, et al. Heme drives hemolysis-induced susceptibility to infection via disruption of phagocyte functions. Nat Immunol. 2016;12:1361–72.

49. Chen H, Lukas TJ, D. N, Suyeoka G, Neufeld AH. Dysfunction of the retinal pigment epithelium with age: increased iron decreases phagocytosis and lysosomal activity. Invest Ophthalmol Vis Sci. 2009;50(4):1895–902.

50. Yefimova MG, Jeanny JC, Keller N, Sergeant C, Guillonneau X, Beaumont C, et al. Impaired retinal iron homeostasis associated with defective phagocytosis in Royal College of Surgeons rats. Invest Ophthalmol Vis Sci. 2002;43(2):537–45.

51. Kimura N, Nakagami G, Minematsu T, Sanada H. Non-invasive detection of local tissue responses to predict pressure ulcer development in mouse models. J Tissue Viability. 2020;29(1):51–7.

52. Karahan A, AAbbasoğlu A, Işık SA, Çevik B, Saltan Ç, Elbaş NÖ, et al. Factors affecting wound healing in individuals with pressure ulcers: A retrospective study. Ostomy/wound management. 2018;64(2):32–9.

53. J.B M Nasser JJ, editors. Redefining Slough: A New Classification System. Symposium on Advanced Wound Care; 2019; Las Vegas, Nevada, USA: HMP Global Learning Network.

54. Nasir NJM, Corrias A, Heemskerk H, Ang ET, Jenkins JH, Sebastin SJ, et al. The panniculus carnosus muscle: a missing link in the chronicity of heel pressure ulcers? J R Soc Interface. 2021;19:1920210631.

55. Kristiansen M, Graversen JH, Jacobsen C, Sonne O, Hoffman HJ, Law SKA, et al. Identification of the haemoglobin scavenger receptor. Nature. 2001;409(6817):198–201.

56. Soares MP, Hamza I. Macrophages and iron metabolism. Immunity. 2016;44(3):492–504.

57. Bessman NJ, Mathieu JRR, Renassia C, Zhou L, Fung TC, Fernandez KC, et al. Dendritic cell-derived hepcidin sequesters iron from the microbiota to promote mucosal healing. Science. 2020;368(6487):186–9.

58. Chen G, Zhang D, Fuchs TA, Manwani D, Wagner DD, Frenette PS. Heme-induced neutrophil extracellular traps contribute to the pathogenesis of sickle cell disease. Blood. 2014;123(24):3818–27.

59. Okubo K, Kurosawa M, Kamiya M, Urano Y, Suzuki A, Yamamoto K, et al. Macrophage extracellular trap formation promoted by platelet activation is a key mediator of rhabdomyolysis-induced acute kidney injury. Nat Med. 2018;24(2):232–8.

60. Ohbuchi A, Kono M, Kitagawa K, Takenokuchi M, Imoto S, Saigo K. Quantitative analysis of hemin-induced neutrophil extracellular trap formation and effects of hydrogen peroxide on this phenomenon. Biochem Biophys Rep. 2017;11:147–53.

61. Perl DP, Good PF. Comparative techniques for determining cellular iron distribution in brain tissues. Ann Neurol. 1992;32:S76–81.

62. Liu S, Grigoryan MM, Vasilevko V, Sumbria RK, Paganini-Hill A, Cribbs DH, et al. Comparative analysis of H&E and Prussian blue staining in a mouse model of cerebral microbleeds. J Histochem Cytochem. 2014;62(11).

63. Cambos M, Scorza T. Robust erythrophagocytosis leads to macrophage apoptosis via a hemin-mediated redox imbalance: role in hemolytic disorders. J Leukoc Biol. 2011;89(1):159–71.

64. Halliwell B. Artefacts with ascorbate and other redox-active compounds in cell culture: epigenetic modifications, and cell killing due to hydrogen peroxide generation in cell culture media. Free Radic Res. 2018;52(9):907–9.

65. Tucker-Kellogg L, Shi Y, White JK, Pervaiz S. Reactive oxygen species (ROS) and sensitization to TRAIL-induced apoptosis, in Bayesian network modelling of HeLa cell response to LY303511. Biochem Pharmacol. 2012;84(10):1307–17.

66. Garry DJ, Ordway GA, Lorenz JN, Radford NB, Chin ER, Grange RW, et al. Mice without myoglobin. Nature. 1998;395(6705):905–8.

67. Gödecke A, Flögel U, Zanger K, Ding Z, Hirchenhain J, Decking UKM, et al. Disruption of myoglobin in mice induces multiple compensatory mechanisms. Proc Natl Acad Sci. 1999;96(18):10495–500.

68. Meeson AP, Radford N, Shelton JM, Mammen PP, DiMaio JM, Hutcheson K, et al. Adaptive mechanisms that preserve cardiac function in mice without myoglobin. Circ Res. 2001;88(7):713–20.

69. Drummen G, van Liebergen L, Op den Kamp J, Post J. C11-BODIPY(581/591), an oxidation-sensitive fluorescent lipid peroxidation probe: (micro)spectroscopic characterization and validation of methodology. Free radical biology & medicine. 2002;33(4):473–90.

70. Ma KL, Wu Y, Zhang Y, Wang GH, Hu ZB, Ruan XZ. Activation of the CXCL16/CXCR6 pathway promotes lipid deposition in fatty livers of apolipoprotein E knockout mice and HepG2 cells. Am J Transl Res. 2018;10(6):1802–16.

71. Anwar A, Keating AK, Joung D, Sather S, Kim GK, Sawczyn KK, et al. Mer tyrosine kinase (MerTK) promotes macrophage survival following exposure to oxidative stress. J Leukoc Biol. 2009;86(1):73–9.

72. Scott RS, McMahon EJ, Pop SM, Reap EA, Caricchio R, Cohen PL, et al. Phagocytosis and clearance of apoptotic cells is mediated by MER. Nature. 2001;10(411):207–11.

73. Tierney MT, Stec MJ, Rulands S, Simons BD, Sacco A. Muscle stem cells exhibit distinct clonal dynamics in response to tissue repair and homeostatic aging. Cell Stem Cell. 2018;22(1):119–27.

74. Schlieper G, Kim JH, Molojavyi A, Jacoby C, Laussmann T, Flogel U, et al. Adaptation of the myoglobin knockout mouse to hypoxic stress. Am J Physiol Regul Integr Comp Physiol. 2004;286(4):R786–92.

75. Hazarika S, Angelo M, Li Y, Aldrich AJ, Odronic SI, Yan Z, et al. Myocyte specific overexpression of myoglobin impairs angiogenesis after hind-limb ischemia. Arterioscler Thromb Vasc Biol. 2008;28(12).

76. Meisner JK, Song J, Annex BH, Price RJ. Myoglobin overexpression inhibits reperfusion in the ischemic mouse hindlimb through impaired angiogenesis but not arteriogenesis. Am J Pathol. 2013;183(6):1710–8.

77. Guardiola O, Andolfi G, Tirone M, Iavarone F, Brunelli S, Minchiotti G. Induction of acute skeletal muscle regeneration by cardiotoxin injection. J Vis Exp. 2017(119):e54515.

78. Harvey AL, Marshall RJ, Karlsson E. Effects of purified cardiotoxins from the Thailand cobra (Naja naja siamensis) on isolated skeletal and cardiac muscle preparations.. Toxicon. 1982;20(2).

79. Averin AS, Utkin YN. Cardiovascular Effects of Snake Toxins: Cardiotoxicity and Cardioprotection. Acta Naturae. 2021;13(3):4–14.

80. Wang Y, Lu J, Liu Y. Skeletal Muscle Regeneration in Cardiotoxin-Induced Muscle Injury Models. Int J Mol Sci. 2022;23(21).

81. Stekelenburg A, Oomens CW, Strijkers GJ, Nicolay K, Bader DL. Compression-induced deep tissue injury examined with magnetic resonance imaging and histology. J Appl Physiol (1985). 2006;100(6):1946–54.

82. Worsley PR, Crielaard H, Oomens CWJ, Bader DL. An evaluation of dermal microcirculatory occlusion under repeated mechanical loads: Implication of lymphatic impairment in pressure ulcers88. Microcirculation. 2020;27(7):e12645.

83. Li J, Cao F, Yin HL, Huang ZJ, Lin ZT, Mao N, et al. Ferroptosis: past, present and future. Cell Death Dis. 2020;11(2):88.

84. Li S, Li Y, Wu Z, Wu Z, Fang H. Diabetic ferroptosis plays an important role in triggering on inflammation in diabetic wound. Am J Physiol Endocrinol Metab. 2021;213(4):E509–E20.

85. Schrijvers DM, De Meyer GR, Kockx MM, Herman AG, Martinet W. Phagocytosis of apoptotic cells by macrophages is impaired in atherosclerosis. Arterioscler Thromb Vasc Biol. 2005;25(6):1256–61.

86. Kao JK, Wang SC, Ho LW, Huang SW, Chang SH, Yang RC, et al. Chronic iron overload results in impaired bacterial killing of THP-1 derived macrophage through the inhibition of lysosomal acidification. PLoS One. 2016;11(5):e0156713.

87. van Asbeck BS, Marx JJM, Struyvenberg A, Verhoef J. Functional defects in phagocytic cells from patients with iron overload. J Infect. 1984;8(3):232–40.

88. Porto G, De Sousa M. Iron overload and immunity. World J Gastroenterol. 2007;13(35):4707–15.

89. Cantinieaux B, Hariga C, Ferster A, de Maertelaere E, Toppet M, Fondu P. Neutrophil dysfunctions in thalassaemia major: The role of cell iron overload. Eur J Haematol. 1987;39(1):28–34.

90. Ali MK, Kim RY, Brown AC, Donovan C, Vanka KS, Mayall JR, et al. Critical role for iron accumulation in the pathogenesis of fibrotic lung disease. J Pathol. 2020;251(1):49–62.

91. Handa P, Thomas S, Morgan-Stevenson V, Maliken BD, Gochanour E, Boukhar S, et al. Iron alters macrophage polarization status and leads to steatohepatitis and fibrogenesis. J Leukoc Biol. 2019;105(5):1015–26.

92. Agoro R, Taleb M, Quesniaux VFJ, Mura C. Cell iron status influences macrophage polarization. PLoS One. 2018;13(5):e0196921.

93. Manna B, Nahirniak P, Morrison CA. Wound Debridement. Treasure Island (FL): StatPearls Publishing; 2021.

94. Eriksson A, Lindström M, Carlsson L, Thornell LE. Hypertrophic muscle fibers with fissures in power-lifters; fiber splitting or defect regeneration?. Histochem Cell Biol. 2006;126(4):409–17.

95. Pichavant C, Burkholder TJ, Pavlath GK. Decrease of myofiber branching via muscle-specific expression of the olfactory receptor mOR23 in dystrophic muscle leads to protection against mechanical stress. Skelet Muscle. 2016;6(2).

96. Soh WSB, Corrias A, Tucker-Kellogg L. Computational modelling of the thin muscle layer, panniculus carnosus, demonstrates principles of pressure injuries and prophylactic dressings. Innovations and Emerging Technologies in Wound Care: Elsevier; 2020. p. 41–52.

97. Ahmed AK, Goodwin CR, Sarabia-Estrada R, Lay F, Ansari AM, Steenbergen C, et al. A non-invasive method to produce pressure ulcers of varying severity in a spinal cord-injured rat model. Spinal Cord. 2016;54(12):1096–104.

98. Siu PM, Tam EW, Teng BT, Pei XM, Ng JW, Benzie IF, et al. Muscle apoptosis is induced in pressure-induced deep tissue injury. J Appl Physiol. 1985;107(4):1266–75.

99. Wassermann E, Van Griensven M, Gstaltner K, Oehlinger W, Schrei K, Redl H. A chronic pressure ulcer model in the nude mouse. Wound Repair Regen. 2009;17(4):480–4.

100. Salcido R, Donofrio JC, Fisher SB, LeGrand EK, Dickey K, Carney JM, et al. Histopathology of pressure ulcers as a result of sequential computer-controlled pressure sessions in a fuzzy rat model. Adv Wound Care. 1994;7(5):23–4.

101. Salcido R, Popescu A, Ahn C. Animal models in pressure ulcer research. J Spinal Cord Med. 2007;30(2):107–16.

102. Cichowitz A, Pan WR, Ashton M. The heel: anatomy, blood supply, and the pathophysiology of PUs. Ann Plast Surg 2009;62(4):423e9.

103. González-Montero J, Brito R, Gajardo AI, Rodrigo R. Myocardial reperfusion injury and oxidative stress: Therapeutic opportunities. World J Cardiol. 2018;10(9):74–86.

104. Morris SF, Pang CY, Lofchy NM, Davidson G, Lindsay WK, Zuker RM, et al. Deferoxamine attenuates ischemia-induced reperfusion injury in the skin and muscle of myocutaneous flaps in the pig. Plast Reconstr Surg. 1993;92(1):120–32.

105. Yang L, Xie P, Wu J, Yu J, Li X, Ma H, et al. Deferoxamine Treatment Combined With Sevoflurane Postconditioning Attenuates Myocardial Ischemia-Reperfusion Injury by Restoring HIF-1/BNIP3-Mediated Mitochondrial Autophagy in GK Rats. Front Pharmacol. 2020;11:6.

106. Peeters-Scholte C, Braun K, Koster J, Kops N, Blomgren K, Buonocore G, et al. Effects of allopurinol and deferoxamine on reperfusion injury of the brain in newborn piglets after neonatal hypoxia-ischemia. Pediatr Res. 2003;54(4):516–22.

107. Qayumi AK, Janusz MT, Dorovini-Zis K, Lyster DM, Jamieson WRE, Poostizadeh A, et al. Additive effect of allopurinol and deferoxamine in the prevention of spinal cord injury caused by aortic crossclamping. J Thorac Cardiovasc Surg. 1994;107(5):1203–9.

108. Zomer HD, Trentin AG. Skin wound healing in humans and mice: Challenges in translational research. J Dermatol Sci. 2018;90(1):3–12.

109. Qin W, Dion SL, Kutny PM, Zhang Y, Cheng AW, Jillette NL, et al. Efficient CRISPR/Cas9-mediated genome editing in mice by zygote electroporation of nuclease. Genetics. 2015;200(2):423–30.

110. Qin W, Kutny PM, Maser RS, Dion SL, Lamont JD, Zhang Y, et al. Generating mouse models using CRISPR-Cas9-mediated genome editing. Curr Protoc Mouse Biol. 2016;6(1):39–66.

111. Barata JD, D’Haese PC, Pires C, Lamberts LV, Simões J, De Broe ME. Low-dose (5 mg/kg) desferrioxamine treatment in acutely aluminium-intoxicated haemodialysis patients using two drug administration schedules. Nephrol Dial Transplant. 1996;11(1):125–32.

112. Shihan MH, Novo SG, Le Marchand SJ, Wang Y, Duncan MK. A simple method for quantitating confocal fluorescent images. Biochemistry and Biophysics Reports. 2021;25:100916.

