## Supplementary Text for "Myoglobin-derived iron causes wound enlargement and impaired regeneration in pressure injuries of muscle"

### Supplementary methods

##### **Pressure wound assessment**

External wounds were assessed daily with a digital calliper (Tactix, CO, USA) positioned at the borders of the wounds to measure the length and width of each wound. In addition, each wound was photographed with an Olympus pocket camera alongside a wound mapping marker (KISS Healthcare Inc., CA, USA). Measurements and photographs were obtained until re-epithelialization (external wound closure). Wound area was computed by ImageJ after manual segmentation of the wound margins.

##### **RAW 264.7 macrophage cell culture and assays**

We cultured RAW 264.7 macrophages (TIB-71, ATCC) in High Glucose DMEM with pyruvate (Gibco, UK) supplemented with 10% Fetal Bovine Serum (FBS; GE Healthcare, USA), 100 units/mL penicillin, 100 µg/mL streptomycin, 0.25 µg/mL Amphotericin B (Fungizone, PSF; GE Healthcare, USA) and 5 µg/mL Plasmocin (GE Healthcare, USA) at 37°C under 5% CO_2_.

RAW 264.7 macrophages were pre-incubated with stimuli for 48 hr before assessment. One stimulus was 20ng/ml recombinant murine IFNγ (Gene Ethics Asia Pte Ltd, Singapore) with 50ng/ml lipopolysaccharide (LPS; L4931), to resemble pro-inflammatory or M1 activation. Another stimulus was 20ng/ml recombinant murine IL-4 (Gene Ethics Asia Pte Ltd, Singapore) to resemble alternately-activated or M2 phenotype. Concentrations and incubation times were based on work by Banete et al. (1).

For the myoglobin stimulus, we used a mouse myoglobin ELISA kit (Abcam, Cambridge, UK, ab157722) to find the myoglobin concentrations in the calf (67.4 µg/ml) and bicep (40.8 µg/ml) muscles of our mice, and found that published literature used similar concentrations (2). From these levels, this we chose the third stimulus to be 50 µg/ml myoglobin (M0630 from horse muscle, Sigma Aldrich, MO, USA). The cells were then measured for viability and cytotoxicity by Cell Titer-Glo^TM^ assay (G9242, Promega, WI, USA) and CytoTox-Glo^TM^ assay (G9292, Promega, WI, USA) respectively.

##### **Protein extraction and detection**

Snap-frozen tissues were homogenized using MAGNAlyser beads (Roche Life Science, Penzberg, Germany) in RIPA lysis buffer (Sigma Aldrich, MO, USA) with protease inhibitors and phosphatase inhibitors (Nacalai Tesque Inc., Kyoto, Japan). The samples were centrifuged for 20 mins and the supernatant homogenate collected. Protein concentrations were determined using the Pierce BCA reagent (Thermofisher, MA, USA).

Luminex ELISA was carried out for the detection of murine interleukins, chemokines and growth factors (R&D Systems, MN, USA; LXSAMSM-21). Sample preparation followed the manufacturer’s protocol, and analytes were read using the MAGPIX® Platform (Luminex Corporation, TX, USA).

For blotting, 30 μg samples were separated by SDS-PAGE (Bio-Rad Laboratories Inc., CA, USA) and transferred to PVDF membranes (Bio-Rad Laboratories Inc., CA, USA). The PVDF membranes were blocked with 5% blotting grade blocker (Bio-Rad Laboratories Inc., CA, USA) at RT for 1 h and then incubated with the primary antibodies (anti-myoglobin antibody, 25919S, Cell Signaling Technology, MA, USA) overnight at 4°C. Then, the membranes were washed with TBST at RT and incubated with the secondary antibody for 1 h. Protein bands were detected with Pierce ECL substrate (Thermofisher, MA, USA) according to the manufacturer's instructions. Data were analyzed with the ChemiDoc Imaging System (Bio-Rad Laboratories Inc., CA, USA).

##### **Iron assay**

An iron assay kit (Abcam, Singapore; ab83366) was used to measure total iron (ferrous (Fe^2+^) and ferric (Fe^3+^)) in the uninjured tibialis anterior (calf) muscles of Mb-/- and Mb+/+ mice. Snap-frozen muscle tissues were homogenized as previously described and the assay was performed according to manufacturer’s instructions.

##### **Confocal Microscopy**

Tissue samples were imaged using a Zeiss LSM710 confocal microscope (Carl Zeiss, Oberkochen, Germany) and Olympus FV3000 laser scanning confocal microscope (Olympus, Tokyo, Japan). Excitation and detection wavelengths used for the respective fluorophores were: CFP^mem^: Ex. 457 nm and Em. 466-495 nm, GFP^nuc^: Ex. 488 nm and Em. 498-510 nm, YFP^cyt^: Ex. 515 nm and Em. 521-560 nm, RFP^cyt^: Ex. 559 nm and Em. 590-650 nm. Images were processed and analysed using Fiji (ImageJ) software. The number of muscle fiber malformations in each mouse was quantified by counting the malformations in a field of view, and averaging over 4 fields of view for each mouse.

##### **Perl’s Prussian blue staining**

8 μm sections of fixed, paraffin-embedded tissues were deparaffinized and rehydrated prior to staining. The sections were then incubated for five minutes in Perl’s Prussian blue iron stain (mix of potassium ferrocyanide and hydrochloric acid; Abcam, Cambridge, UK, ab150674) rinsed in water, and counterstained in nuclear fast red (Abcam, Singapore, ab150674). Iron deposits were observed as deep blue dots or speckles, and the number and fractional area occupied by deep blue pixels were quantified by thresholding in Matlab (Mathworks, MA, USA).

##### **BODIPY 581/591 C11 detection**

10 μm tissue cryosections were fixed in 4% paraformaldehyde, blocked with 10% normal serum and permeabilized with 0.2% Tween 20. Tissue sections were stained with 10 μM BODIPY 581/591 C11 probe (Thermofisher, MA, USA, D3861). After 30 minutes in the dark, the slides were mounted with Vectashield Hardset with DAPI (Vector Laboratories, CA, USA) and viewed under a Leica TCS SP8 confocal microscope (Leica Microsystems, Wetzlar, Germany). Each wound section was imaged from edge to edge with consecutive frames.

References

1.	Banete A ea. Immortalized murine macrophage cell line as a model for macrophage polarization into classically activated M(IFNγ+LPS) or alternatively activated M(IL-4) macrophages. J Clin Cell Immunol. 2015;6(2).

2.	Belliere J ea. Specific macrophage subtypes influence the progression of rhabdomyolysis-induced kidney injury. J Am Soc Nephrol. 2015;26(6):1363-77.
