## Supplementary Figures & Tables for "Myoglobin-derived iron causes wound enlargement and impaired regeneration in pressure injuries of muscle"

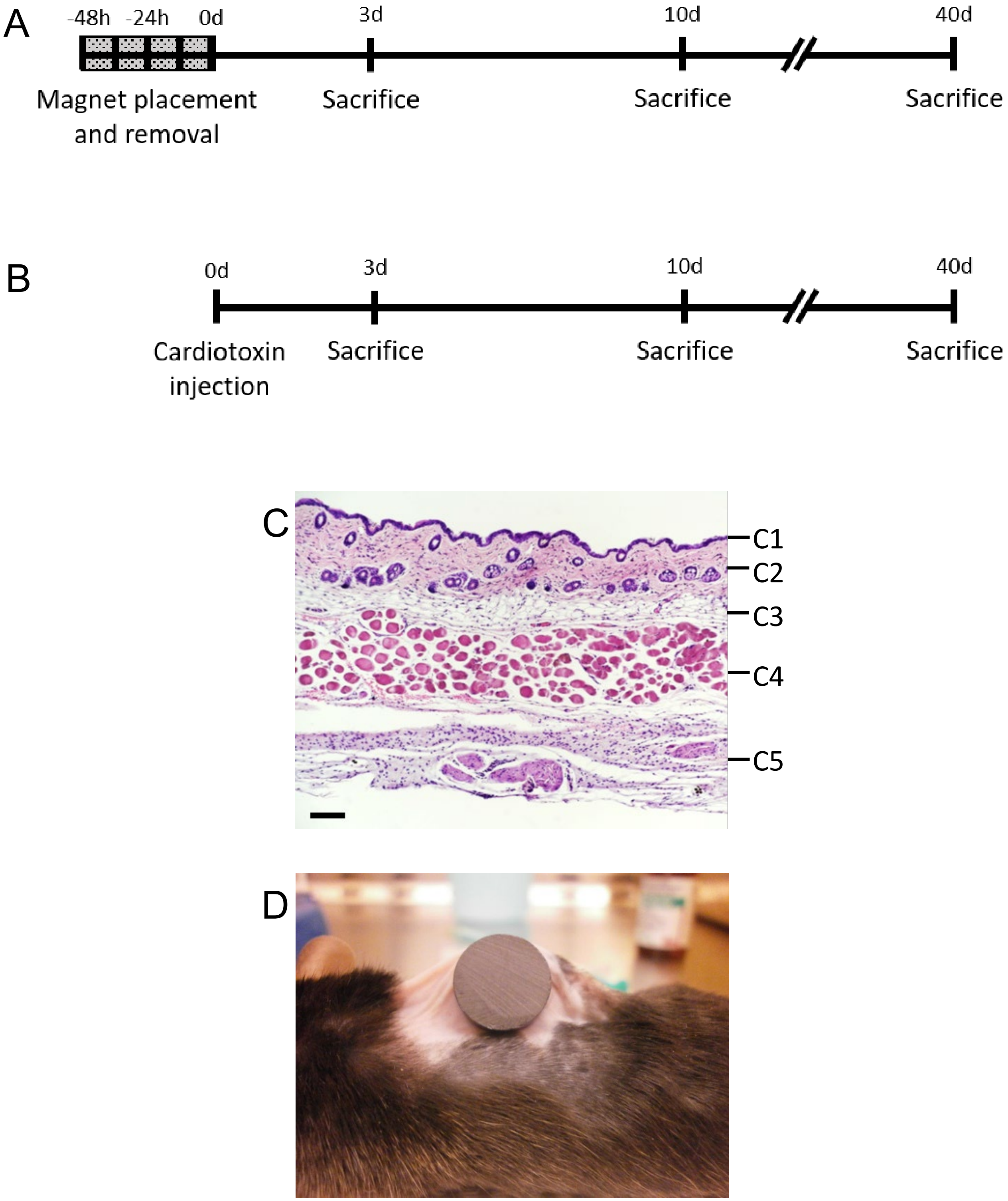


**Supplementary Figure 1: Mouse model of injury.** (A) Experimental schedule of mPI induction and tissue collection. (B) Experimental schedule of cardiotoxin acute injury and tissue collection. (C) An H&E stained cross-section of healthy murine dorsal skinfold, showing the epidermis (C1), dermis (C2), dermal white adipose tissue, dWAT (C3), *panniculus carnosus* muscle, PC (C4), and loose areolar tissue (C5). Scale bar is 50 µm. (D) Image of mPI induction in C57BL6 mouse with a 12 mm magnet on the dorsal skinfold.

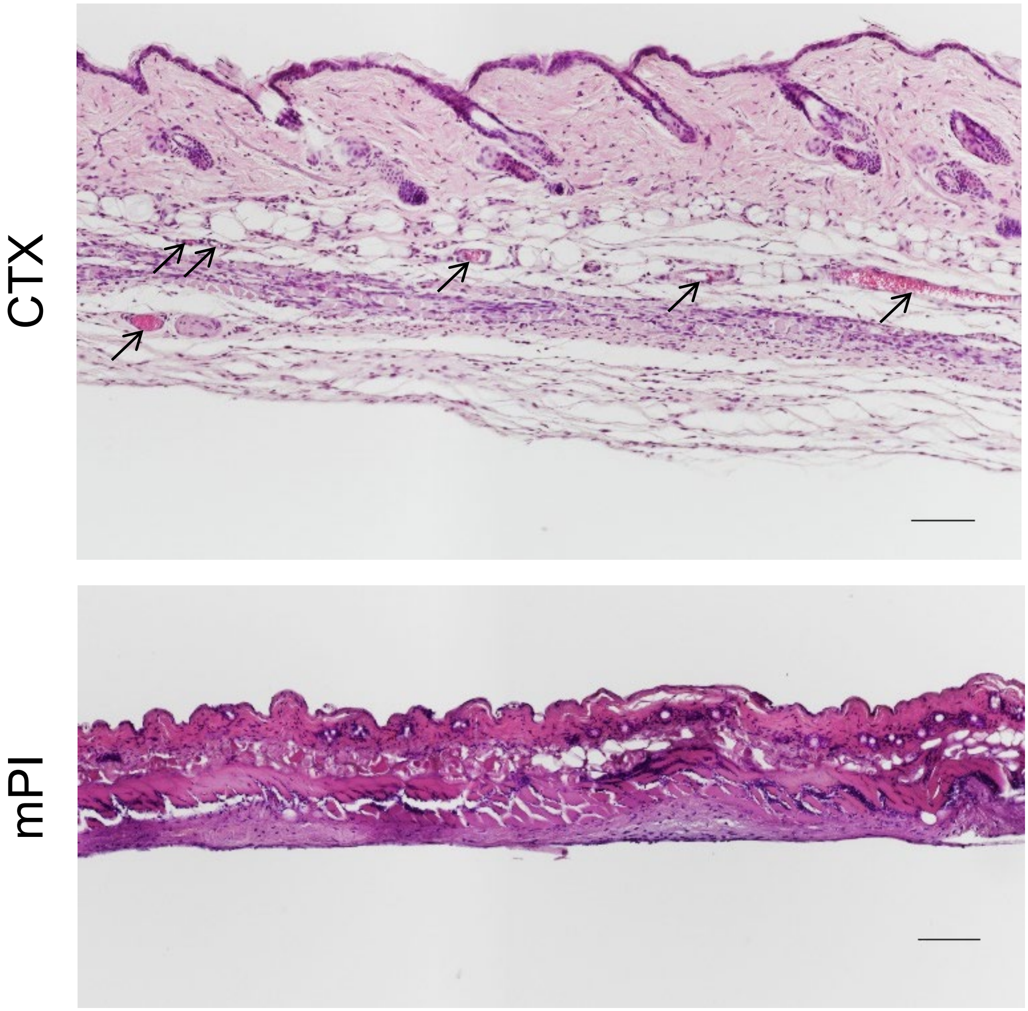


**Supplementary Figure 2: Intact blood vessels are absent or infrequent in the dead tissue of muscle pressure injuries (mPI).** H&E sections of CTX-injured tissue (top) versus mPI (bottom) at day 3. Black arrows point to intact capillaries in the skinfold. Scale bars: 100 µm.

| Wound healing milestones | Cardiotoxin (Acute injury) | | | | Pressure injury (Chronic wound) | | | |
|---|---|---|---|---|---|---|---|---|
| | 3 Days | 10 Days | 40 Days | 90 Days | 3 Days | 10 Days | 40 Days | 90 Days |
| Immune cells pervade the wounded area | ✓ | | | | No | ✓ | | |
| Dead tissue fully cleared | No | ✓ | | | No | No | ✓ | |
| Wounded area revascularized | No | ✓ | | | No | No | ✓ | |
| Immature myotubes have begun to form | No | ✓ | | | No | No | ✓ | |
| Immature myotubes fill the wounded area | No | ✓ | | | No | No | No | No |
| Mature muscle fibers fill the wounded area | No | No | ✓ | | No | No | No | No |

**Supplementary Figure 3: Schematic** showing differences in injury response and regeneration between cardiotoxin (CTX, acute injury) and muscle pressure injury (mPI, chronic wound).

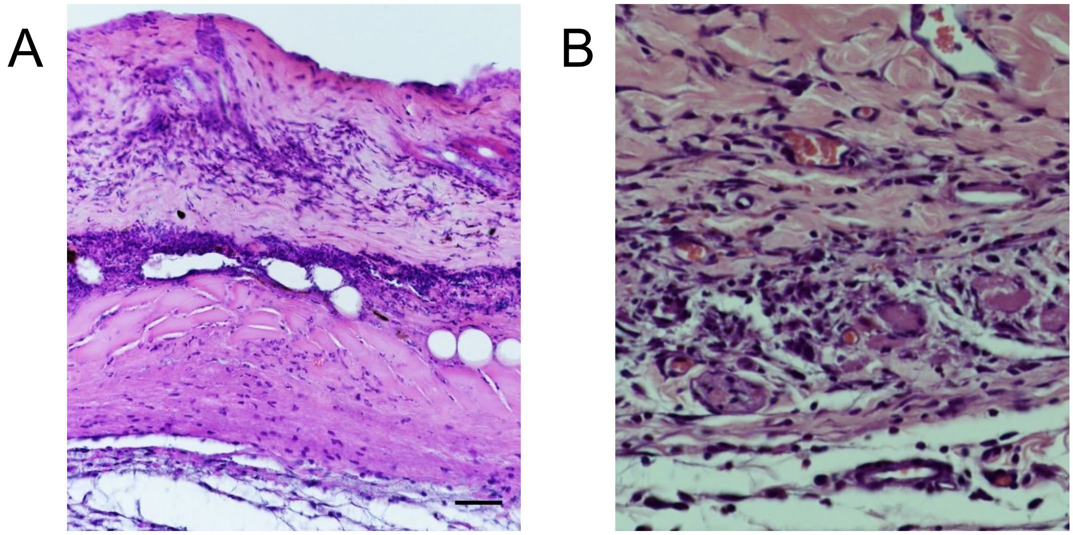


**Supplementary Figure 4: Wound margins show intact immune cells but the compressed region of mPI lack viable immune infiltrate.** (A) A cross-section of mPI at day 3 post-injury, stained with H&E showing indicators of cell death (karyolysis, karyorrhexis and acidification), and a lack of intact immune cells in the compressed region. (B) Another cross-section taken from the wound edge (boundary between injured and uninjured tissue) showing intact immune cell infiltrate. Scale bar is 25 μm.

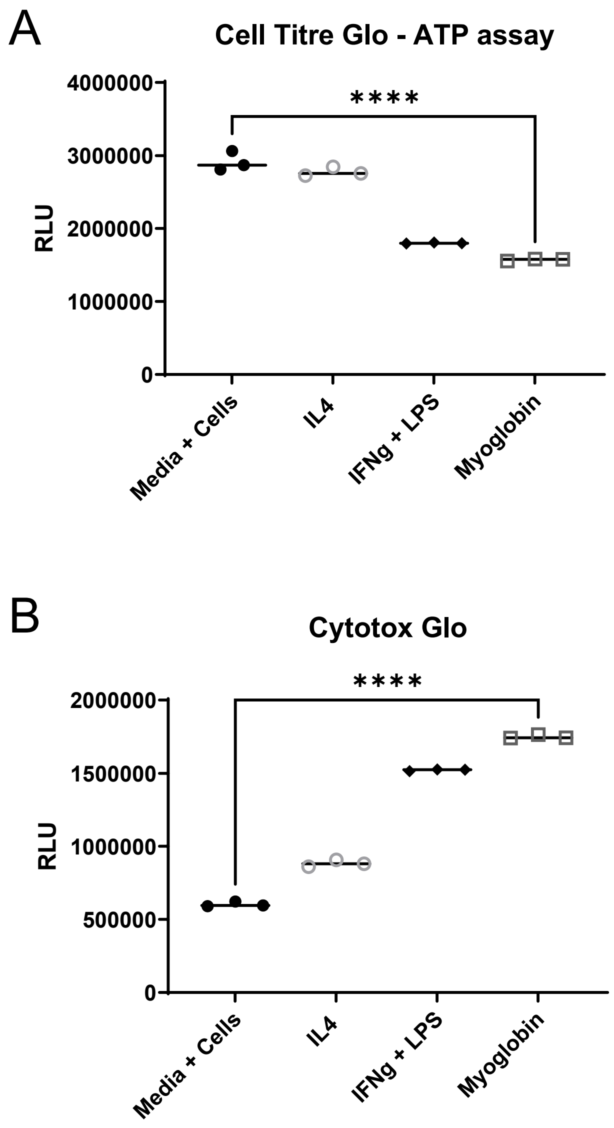


**Supplementary Figure 5: Extracellular myoglobin decreases the viability of monocytic cells *in vitro*.** RAW 264.7 monocyte/macrophage-like cells were incubated with myoglobin at 50 µg/mL, the level in muscle lysate, or with traditional stimuli for pro-inflammatory (20ng/ml IFNγ + 50ng/ml LPS) or alternately-activated (20ng/ml IL4) states for 48 hours prior to assessment. (A) A viability measure (Cell Titre-Glo assay for ATP) (b) a cytotoxicity measure (Cytotox-Glo assay for release of dead cell protease) of cells in the various treatment groups.

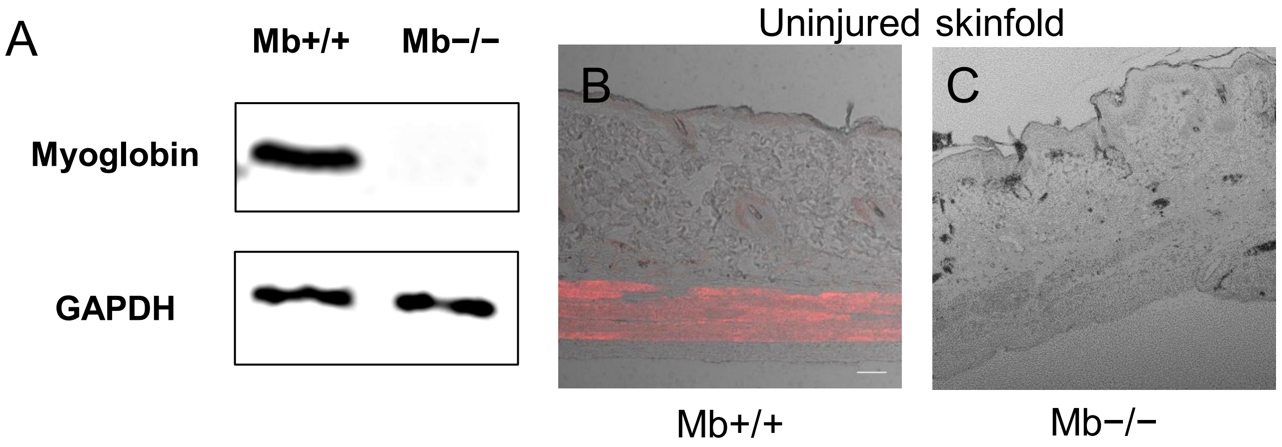


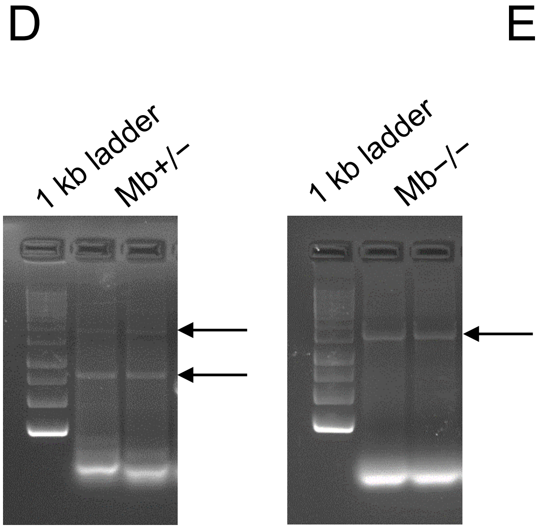


E

| Tissue layer | Measurement (μm) | | p-value |
|---|---|---|---|
| | Mb+/+ | Mb−/− | |
| Epidermis | 19.6 ± 4.70 | 17.3 ± 3.90 | 0.209 |
| Dermis | 305 ± 42.4 | 272 ± 52.8 | 0.117 |
| dWAT | 107 ± 28.7 | 157 ± 47.5 | 0.023 (*) |
| Panniculus carnosus | 112 ± 15.4 | 120 ± 37.4 | 0.545 |
| Loose areolar tissue | 71.3 ± 18.9 | 80.8 ± 25.2 | 0.391 |
| Capillary density | 100% | 117% | 0.046 (*) |

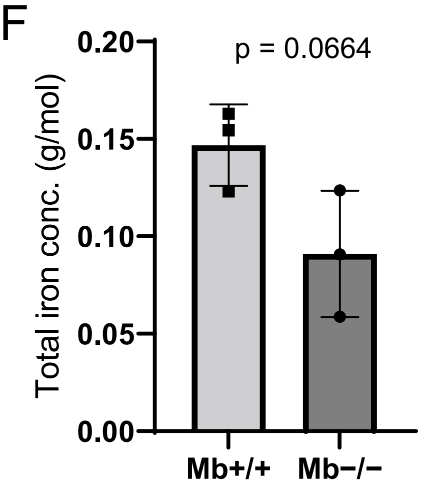


G

| 2-year-old mice (male C57BL6) | Treatment | Endpoint |
|---|---|---|
| Myoglobin+/+ (n = 3) | Saline | 3 days |
| Myoglobin−/− (n = 3) | Saline | 3 days |

**Supplementary Figure 6: Validation of Myoglobin-knockout mice.** (A) Western blotting in panniculus carnosus homogenate to validate the myoglobin levels of Mb+/+ and Mb−/− mice (one blot for every mouse). (B-C) Myoglobin immuno-staining (red) in the uninjured skinfold to validate the myoglobin levels of Mb+/+ and Mb−/− mice (at least four samples per mouse). Scale bars are 50 μm. (D) Gel electrophoresis bands from PCR amplification of Mb−/− (1700 bp; top arrows) and Mb+/+ (648 bp; bottom arrow) from mouse genomic DNA. (E) Morphological comparison of the dorsal skinfolds (measured from H&E sections) in Mb+/+ and Mb−/− mice. (F) Quantification of total iron in *tibialis anterior* muscles in Mb+/+ and Mb−/− mice. (G) Table of treatment arms.

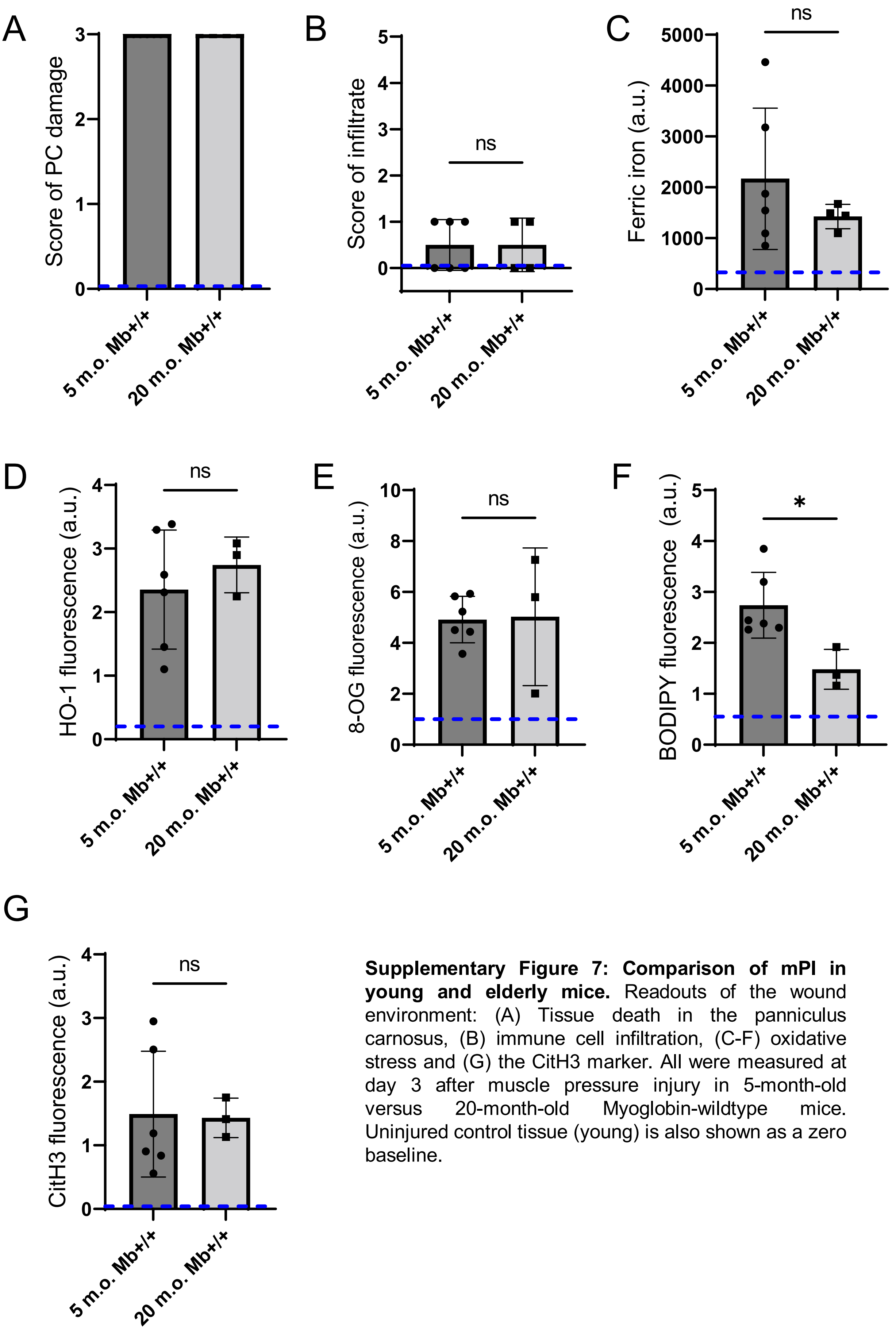


**Supplementary Figure 7: Comparison of mPI in young and elderly mice.** Readouts of the wound environment: (A) Tissue death in the panniculus carnosus, (B) immune cell infiltration, (C-F) oxidative stress and (G) the CitH3 marker. All were measured at day 3 after muscle pressure injury in 5-month-old versus 20-month-old Myoglobin-wildtype mice. Uninjured control tissue (young) is also shown as a zero baseline.

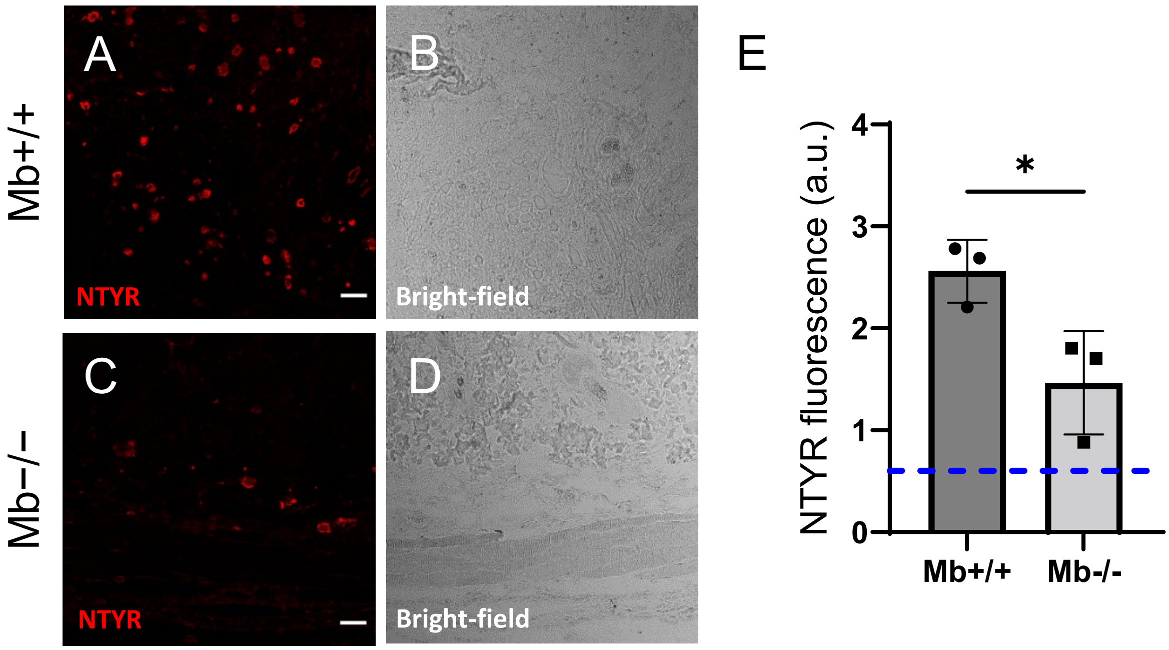


**Supplementary Figure 8: Nitroxidative stress in Mb+/+ versus Mb−/− tissues three days after mPI.** (A-D) Immunostaining of nitrotyrosine (NTYR), a marker of nitroxidative stress, in Mb+/+ versus Mb−/− pressure-injured tissues. (B) and (D) are bright-field images of (A) and (C), respectively. Scale bars are 50 μm. (E) Quantification of NTYR staining. Blue dotted line indicates average NTYR fluorescence in uninjured skinfold.

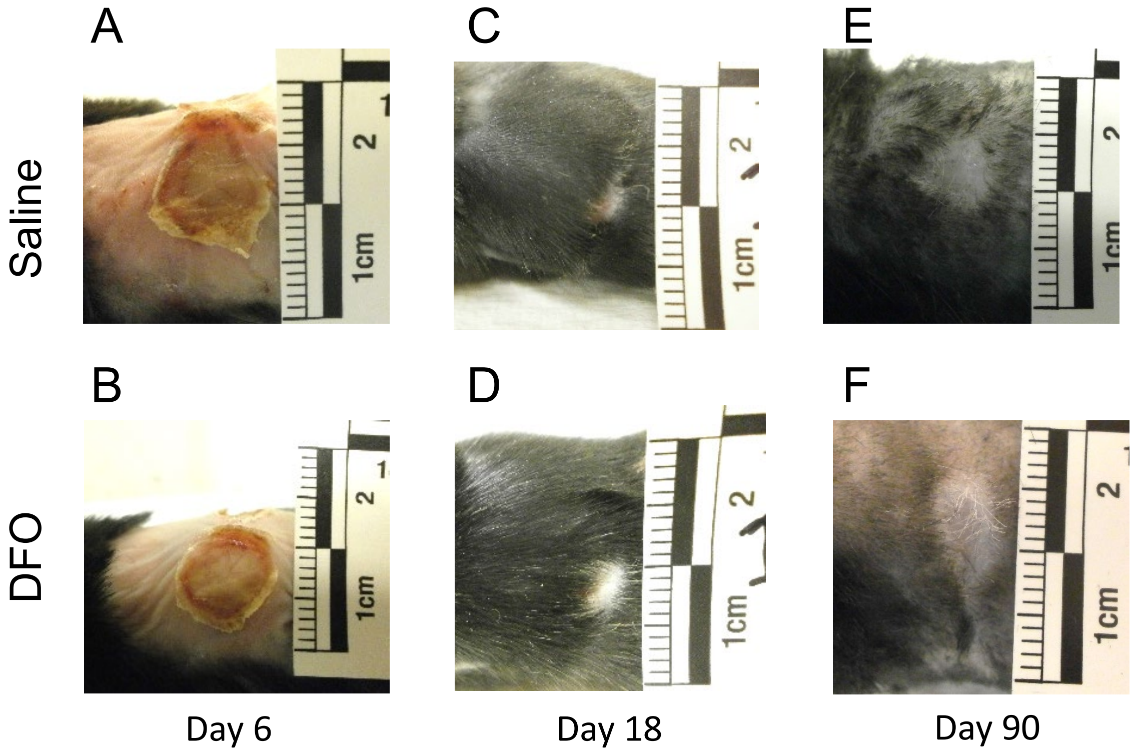


**Supplementary Figure 9: DFO treatment did not significantly affect external wound sizes nor times to wound closure.** (A-F) Photographs of the external wounds of saline- versus DFO-treated mice at (A-B) six, (C-D) 18 and (E-F) 90 days after injury.

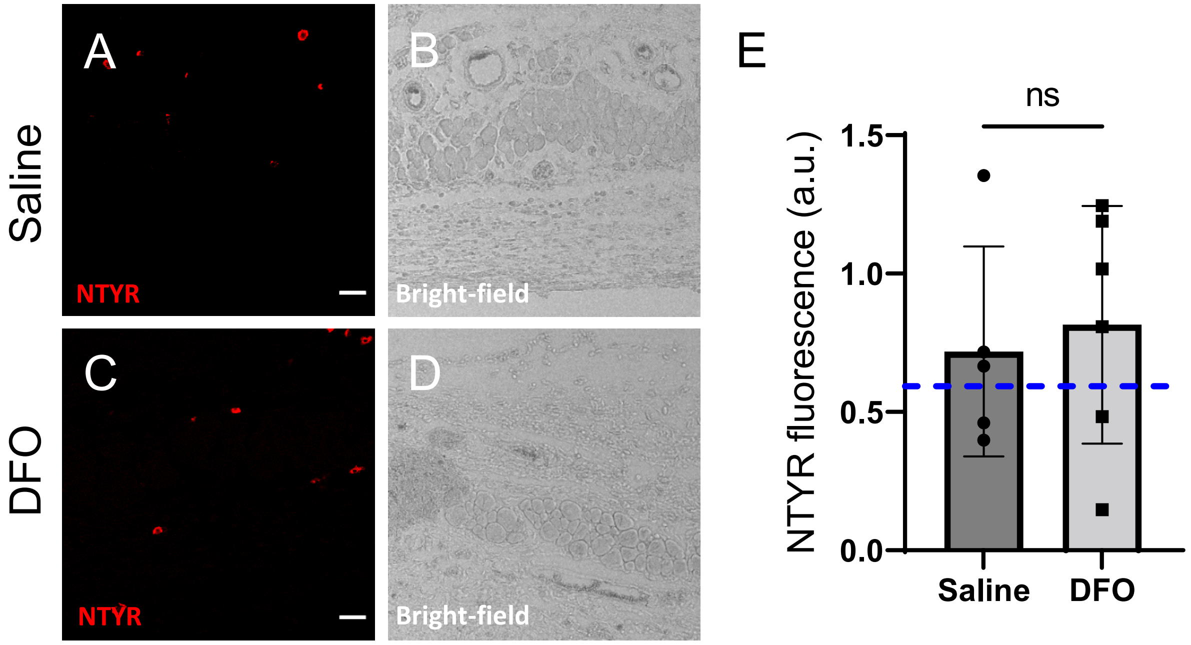


**Supplementary Figure 10: Nitroxidative stress in DFO versus saline-treated tissues three days after mPI.** (A-D) Immunostaining of nitrotyrosine (NTYR), a marker of protein nitration and nitroxidative stress, in saline- versus DFO-treated wound tissues. (B) and (D) are bright-field images of (A) and (C), respectively. Scale bars are 50 µm. (E) Quantification of NTYR staining. Blue dotted line indicates average NTYR fluorescence in uninjured skinfold.

Saline

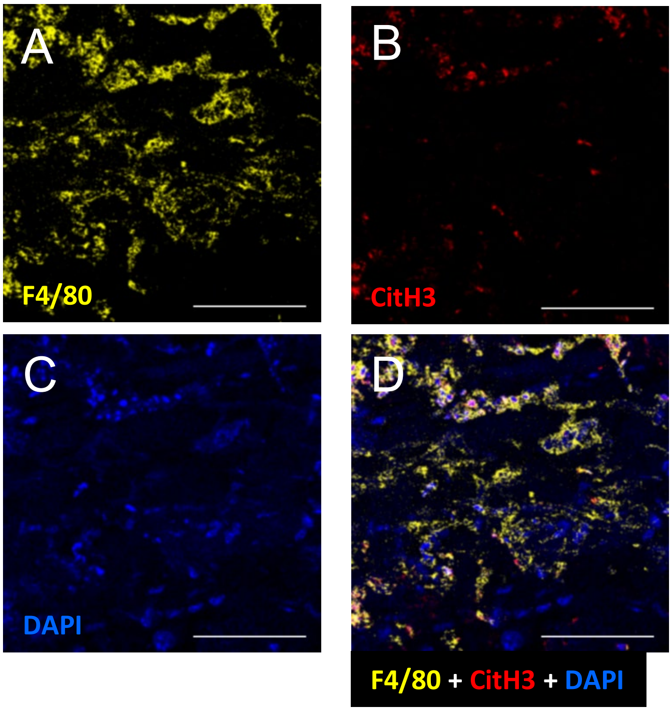


E

Saline

F

DFO

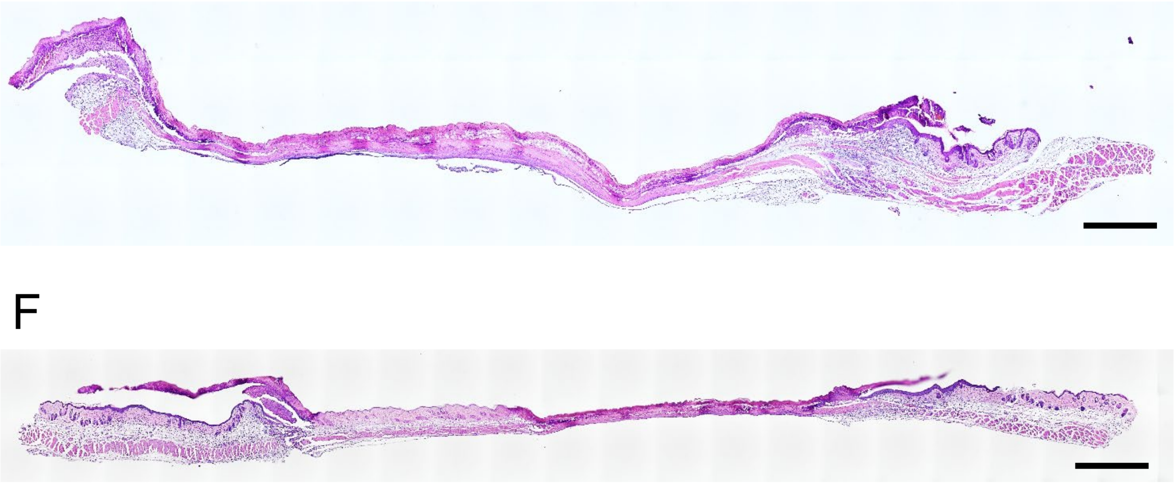

G

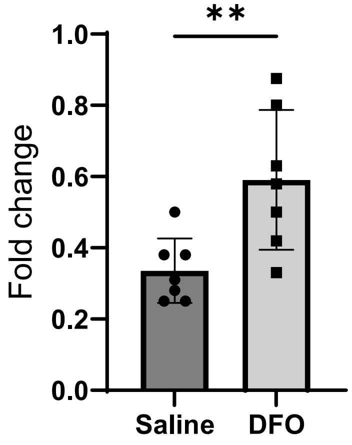


**Supplementary Figure 11: Injury response at day 3 after pressure.** (A) Immuno-staining in yellow for F4/80 receptor (a pan-macrophage marker), (B) CitH3, a marker of extracellular traps, in red and (C) DAPI nuclear stain. (D) Merge of (A), (B) and (C). All images were collected from saline-treated mPI at day 3 post-injury. Laser excitations were sequential, not simultaneous, using 555 nm for yellow, and 594 nm for red. Scale bars are 50 µm. H&E-stained wound sections of (E) saline- and (F) DFO-treated mPI at 3 days. Scale bars are 1000 µm. (G) Another difference between DFO and saline at day 3 was the thickness of the PC at the edge of the wound. This might reflect edema.

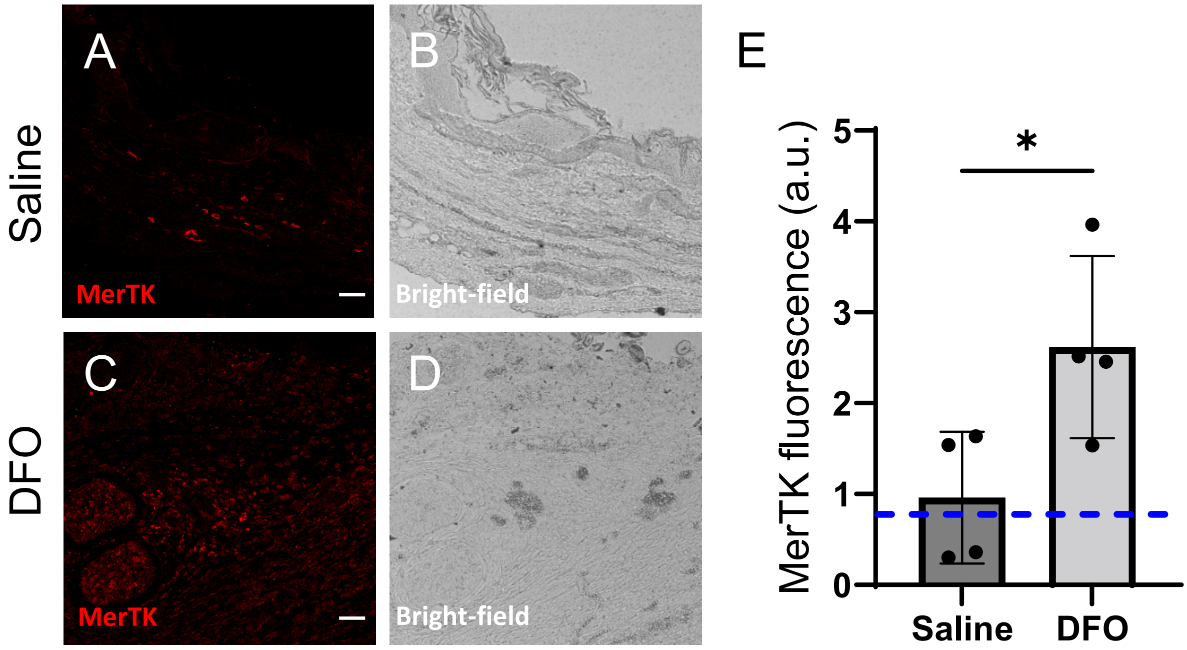


**Supplementary Figure 12: Immune infiltration and function in DFO versus saline-treated tissues seven days after mPI.** (A-D) Immunostaining of MerTK, a marker of macrophage phagocytosis, in saline- versus DFO-treated wound tissues. (B) and (D) are bright-field images of (A) and (C), respectively. Scale bars are 50 μm. (E) Quantification of MerTK staining.

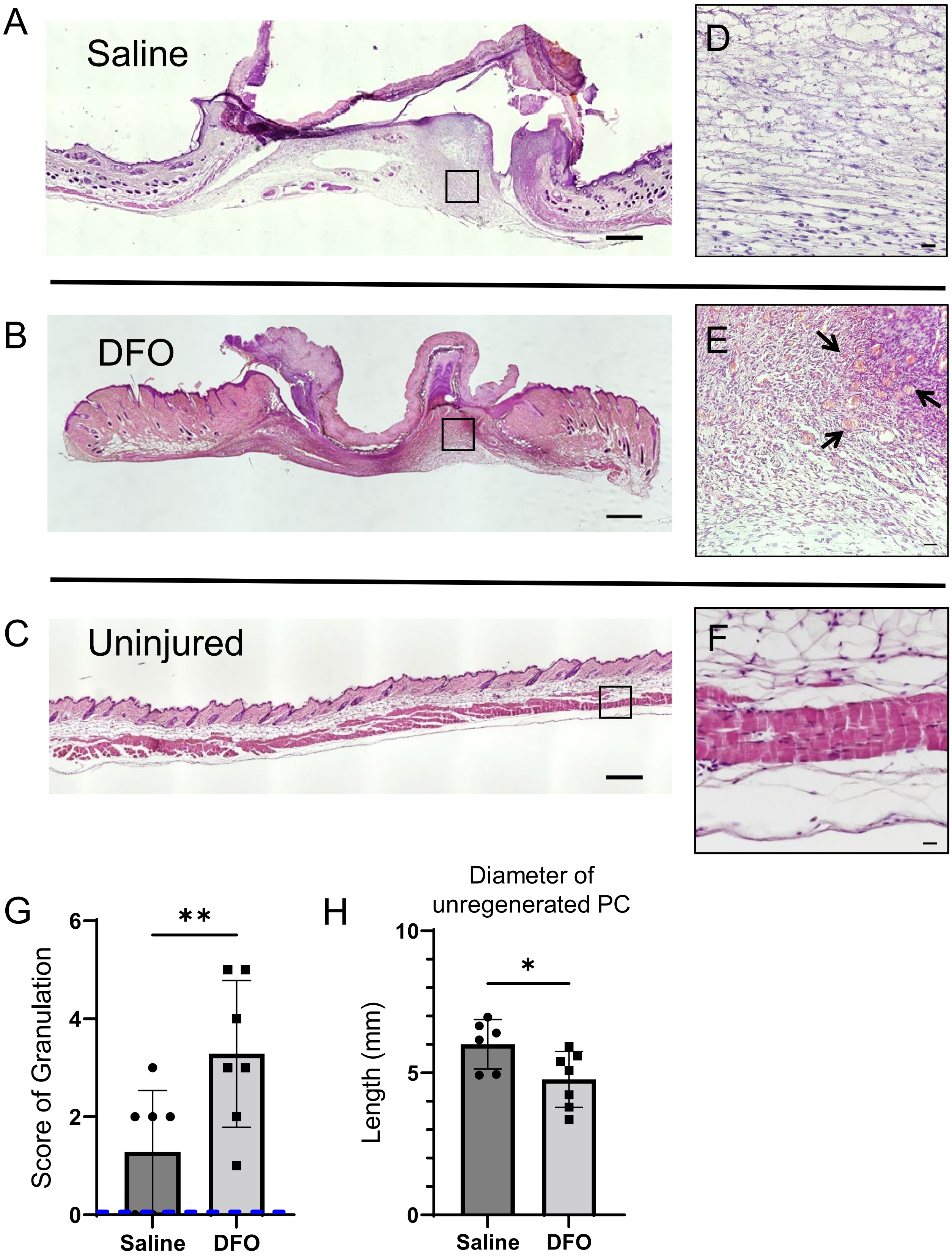


**Supplementary Figure 13: DFO treatment improved angiogenesis and granulation following mPI.** Cross-sections of (A) saline-treated and (B) DFO-treated mPI with surrounding margins, at 10 days post-injury, stained with H&E. (C) Cross-section of healthy uninjured skinfold, stained with H&E. Scale bars are 500 μm. (D-F) Close-ups of (A-C) respectively. Arrows point to new capillaries of granulation tissue. Scale bars are 10 μm. (G) Histopathology scoring of granulation in saline- versus DFO-treated mPI at day 10. (H) Diameter of the unregenerated PC (diameter of hole) of DFO-treated tissues versus saline 10 days after mPI.

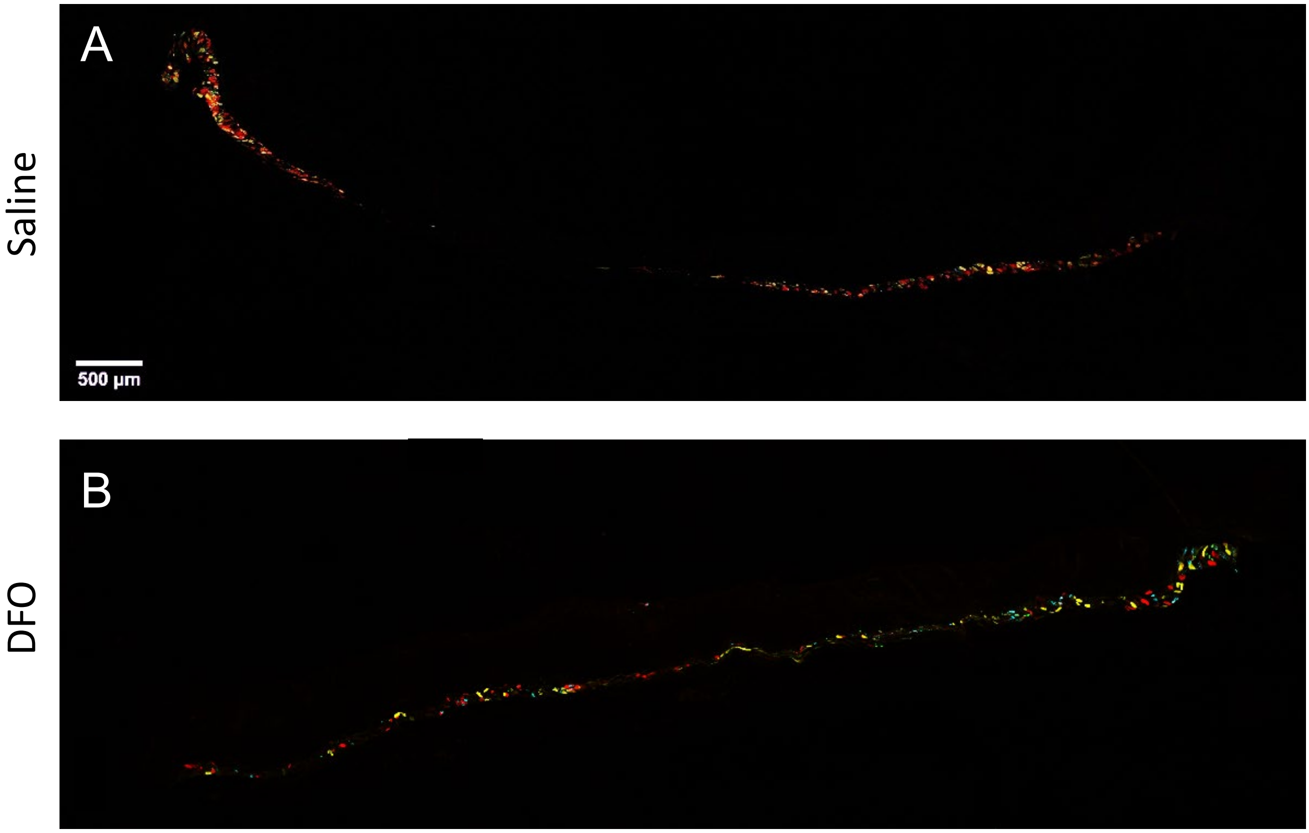


**Supplementary Figure 14: Failure of muscle regeneration after mPI is ameliorated by DFO treatment.** (A-B) Confocal fluorescent imaging of wound cross-sections from (A) saline- and (B) DFO-treated samples, 40 days post-injury, showing the presence or absence of regenerated myofibers via confetti fluorescence in the *panniculus carnosus* muscle.

| Time-point | Cardiotoxin injury, diameter of dead muscle (mm) | Muscle pressure injury, diameter of dead muscle (mm) | p-value |
|---|---|---|---|
| Day 3 | 8.44 ± 0.89 | 8.23 ± 0.98 | 0.5782 (ns) |
| Day 10 | 0.00 ± 0<br>(muscle has regenerated) | 5.75 ± 0.97 | < 0.0001 (****) |

**Supplementary Table 1:** Injuries to the panniculus carnosus muscle from CTX and mPI have comparable diameters at day 3, but are significantly different at day 10.

| Time-point | External Wound Area | |
|---|---|---|
| | **Myoglobin+/+** | **Myoglobin−/−** |
| Day 0 | 0.0 $cm^2$ | 0.0 $cm^2$ |
| Day 1 | 0.290 ± 0.069 $cm^2$ | 0.224 ± 0.016 $cm^2$ |
| Day 2 | 0.254 ± 0.050 $cm^2$ | 0.165 ± 0.034 $cm^2$ |
| Day 3 | 0.219 ± 0.0314 $cm^2$ | 0.105 ± 0.083 $cm^2$ |

**Supplementary Table 2:** External wound area in Myoglobin−/− and age- and sex-matched Myoglobin+/+ mice in the initial days following mPI, using 5 mm magnets. n = 2.

| Analyte | Measurement | | p-value |
|---|---|---|---|
| | Myoglobin+/+ | Myoglobin−/− | |
| CCL3 | 0.398 ± 0.312 pg/ml | 0.364 ± 0.096 pg/ml | 0.889 (ns) |
| CCL5 | 39.3 ± 3.13 pg/ml | 42.5 ± 10.1 pg/ml | 0.690 (ns) |
| CCL7 | 1.5786 ± 0.0340 pg/ml | 2.58 ± 0.512 pg/ml | 0.051 |
| CXCL12 | 296.62 ± 109.45 pg/ml | 150 ± 24.7 pg/ml | 0.138 (ns) |
| CXCL16 | 12.6 ± 2.85 pg/ml | 3.55 ± 0.54 pg/ml | 0.011 (*) |
| IL1b | 162.11 ± 164.06 pg/ml | 232 ± 144 pg/ml | 0.673 (ns) |
| IL4 | 21.616 ± 8.311 pg/ml | 21.0 ± 5.08 pg/ml | 0.937 (ns) |
| IL6 | 14.026 ± 1.591 pg/ml | 15.5 ± 1.54 pg/ml | 0.410 (ns) |
| IL10 | 4.5928 ± 0.0456 pg/ml | 4.66 ± 0.237 pg/ml | 0.722 (ns) |
| RAGE/AGER | 52.95 ± 47.448 pg/ml | 88.5 ± 23.2 pg/ml | 0.395 (ns) |
| uPAR | 262.47 ± 43.09 pg/ml | 485 ± 168 pg/ml | 0.144 (ns) |
| VEGF | 64.441 ± 3.711 pg/ml | 56.1 ± 19.3 pg/ml | 0.581 (ns) |
| PDGF-AA | 27.247 ± 6.276 pg/ml | 17.2 ± 7.48 pg/ml | 0.219 (ns) |
| PAI-1 | 22.5 ± 2.68 pg/ml | 36.9 ± 3.12 pg/ml | 0.008 (**) |
| IGF1 | 28.5 ± 3.44 pg/ml | 28.0 ± 4.97 pg/ml | 0.902 (ns) |
| Endoglin | 90.3 ± 82.4 pg/ml | 2406 ± 3355 pg/ml | 0.384 (ns) |

**Supplementary Table 3:** Luminex measures of various cytokines, chemokines and growth factors between Myoglobin+/+ and Myoglobin−/− tissues, three days after mPI. The p values were computed using a Student's t-test with Bonferroni-Dunn correction.

| Treatment | Endpoint | # of mice |
|---|---|---|
| Saline control | 3 days | n = 7 |
| | 7 days | n = 4 |
| | 10 days | n = 7 |
| | 40 days | n = 7 |
| | 90 days | n = 5 |
| DFO | 3 days | n = 7 |
| | 7 days | n = 4 |
| | 10 days | n = 7 |
| | 40 days | n = 7 |
| | 90 days | n = 5 |

**Supplementary Table 4: Treatment arms for 5-month-old mice with mPI.** Both sexes were used. Mice were sex-matched and age-matched, and littermate controls were chosen when available.

| Analyte | Measurement | | p-value |
|---|---|---|---|
| | Saline-treated | DFO-treated | |
| CCL3 | 1680 ± 275 pg/ml | 1360 ± 539 pg/ml | 0.207 (ns) |
| CCL5 | 166 ± 102 pg/ml | 117 ± 47.0 pg/ml | 0.312 (ns) |
| CCL7 | 258 ± 260 pg/ml | 280 ± 276 pg/ml | 0.892 (ns) |
| CXCL12 | 3500 ± 3590 pg/ml | 3070 ± 4310 pg/ml | 0.854 (ns) |
| CXCL16 | 200 ± 45.6 pg/ml | 128 ± 39.0 pg/ml | 0.017 (*) |
| IL1b | 625 ± 662 pg/ml | 299 ± 177 pg/ml | 0.267 (ns) |
| IL4 | 44.1 ± 32.2 pg/ml | 26.9 ± 10.6 pg/ml | 0.003 (**) |
| IL6 | 95.6 ± 67.5 pg/ml | 77.4 ± 30.5 pg/ml | 0.558 (ns) |
| IL10 | 30.7 ± 27.8 pg/ml | 9.18 ± 3.97 pg/ml | 0.085 (ns) |
| RAGE/AGER | 128 ± 172 pg/ml | 0.00 pg/ml | 0.093 (ns) |
| uPAR | 12800 ± 9739 pg/ml | 6840 ± 3080 pg/ml | 0.179 (ns) |
| VEGF | 1060 ± 457 pg/ml | 981 ± 431 pg/ml | 0.761 (ns) |
| PDGF-AA | 209 ± 85.5 pg/ml | 165 ± 81.3 pg/ml | 0.378 (ns) |
| PAI-1 | 7850 ± 2140 pg/ml | 11400 ± 9550 pg/ml | 0.386 (ns) |
| IGF1 | 2540 ± 834 pg/ml | 2540 ± 1600 pg/ml | 0.996 (ns) |
| Endoglin | 41600 ± 27500 pg/ml | 15600 ± 14400 pg/ml | 0.063 (ns) |

**Supplementary Table 5:** Luminex measures of various cytokines, chemokines and growth factors between saline- and DFO-treated, three days after mPI. The p values were computed using a Student's t-test with Bonferroni-Dunn correction.

| Analyte | Measurement | | p-value |
|---|---|---|---|
| | Saline-treated | DFO-treated | |
| CCL3 | 1880 ± 1060 pg/ml | 1680 ± 855 pg/ml | 0.743 (ns) |
| CCL5 | 230 ± 143 pg/ml | 168 ± 67.8 pg/ml | 0.508 (ns) |
| CCL7 | 557 ± 479 pg/ml | 303 ± 228 pg/ml | 0.363 (ns) |
| CXCL12 | 8920 ± 9850 pg/ml | 3720 ± 3240 pg/ml | 0.251 (ns) |
| CXCL16 | 315 ± 181 pg/ml | 185 ± 50 pg/ml | 0.101 (ns) |
| IL1b | 726 ± 408 pg/ml | 569 ± 349 pg/ml | 0.507 (ns) |
| IL4 | 57.1 ± 13.8 pg/ml | 46.3 ± 18.9 pg/ml | 0.309 (ns) |
| IL6 | 317 ± 283 pg/ml | 160 ± 103 pg/ml | 0.236 (ns) |
| IL10 | 33.9 ± 25.7 pg/ml | 48.6 ± 42.3 pg/ml | 0.508 (ns) |
| RAGE/AGER | 436 ± 536 pg/ml | 268 ± 277 pg/ml | 0.913 (ns) |
| uPAR | 13400 ± 9250 pg/ml | 14100 ± 6060 pg/ml | 0.889 (ns) |
| VEGF | 365 ± 485 pg/ml | 2480 ± 2130 pg/ml | 0.324 (ns) |
| PDGF-AA | 172 ± 111 pg/ml | 213 ± 141 pg/ml | 0.602 (ns) |
| PAI-1 | 23500 ± 36300 pg/ml | 14900 ± 5750 pg/ml | 0.425 (ns) |
| IGF1 | 5670 ± 4800 pg/ml | 3120 ± 1440 pg/ml | 0.244 (ns) |
| Endoglin | 38800 ± 22400 pg/ml | 44800 ± 21100 pg/ml | 0.657 (ns) |

**Supplementary Table 6:** Luminex measures of various cytokines, chemokines and growth factors between saline- and DFO-treated, ten days after mPI. The p values were computed using a Student's t-test with Bonferroni-Dunn correction.

**Health Report**

Duke-NUS Vivarium

| Sentinel Strain: ICR<br>Room: SPF room | | | Species: Mice | | | | Health Status: SPF | | |
|---|---|---|---|---|---|---|---|---|---|
| **Test Reference:** | **7341-2019** | **22559-2019** | **33873-2019** | **9872-2020** | **24231-2020** | **33654-2020** | **11511-2-21** | **Test Laboratory** | **Test Methods** |
| **Date tested:** | **6/3/2019** | **7/8/2019** | **6/12/2019** | **30/3/2020** | **23/9/2020** | **18/12/2020** | **7/4/2021** | | |
| **SEROLOGY (QUARTERLY)** | | | | | | | | | |
| Ectromelia Virus (Mousepox) | 0/5 | 0/5 | 0/5 | 0/6 | 0/6 | 0/6 | 0/6 | RADIL | PCR/MFI |
| Epizootic diarrhoea of Infant Mice (EDIM) | 0/5 | 0/5 | 0/5 | 0/6 | 0/6 | 0/6 | 0/6 | RADIL | PCR/MFI |
| Lymphocytic Choriomeningitis Virus (LCMV) | 0/5 | 0/5 | 0/5 | 0/6 | 0/6 | 0/6 | 0/6 | RADIL | PCR/MFI |
| Mycoplasma pulmonis | 0/5 | 0/5 | 0/5 | 0/6 | 0/6 | 0/6 | 0/6 | RADIL | PCR/MFI |
| Mouse Hepatitis Virus (MHV) | 0/5 | 0/5 | 0/5 | 0/6 | 0/6 | 0/6 | 0/6 | RADIL | PCR/MFI |
| Murine norovirus (MNV) | 0/5 | 0/5 | 0/5 | 0/6 | 0/6 | 0/6 | 0/6 | RADIL | PCR/MFI |
| Mouse parvovirus (MPV) | 0/5 | 0/5 | 0/5 | 0/6 | 0/6 | 0/6 | 0/6 | RADIL | PCR/MFI |
| Minute Virus of Mice (MVM) | 0/5 | 0/5 | 0/5 | 0/6 | 0/6 | 0/6 | 0/6 | RADIL | PCR/MFI |
| Pneumonia virus of mice (PVM) | 0/5 | 0/5 | 0/5 | 0/6 | 0/6 | 0/6 | 0/6 | RADIL | PCR/MFI |
| Mouse Reovirus Type 3 | 0/5 | 0/5 | 0/5 | 0/6 | 0/6 | 0/6 | 0/6 | RADIL | PCR/MFI |
| Theiler's Murine Encephalomyelitis Virus | 0/5 | 0/5 | 0/5 | 0/6 | 0/6 | 0/6 | 0/6 | RADIL | PCR/MFI |
| Sendai Virus | 0/5 | 0/5 | 0/5 | 0/6 | 0/6 | 0/6 | 0/6 | RADIL | PCR/MFI |
| CAR bacillus | 0/5 | - | - | 0/6 | - | - | 0/6 | RADIL | PCR/MFI |
| Encephalitozoon cuniculi | 0/5 | - | - | 0/6 | - | - | 0/6 | RADIL | PCR/MFI |
| MAD1 | 0/5 | - | - | 0/6 | - | - | 0/6 | RADIL | PCR/MFI |
| MAD2 | 0/5 | - | - | 0/6 | - | - | 0/6 | RADIL | PCR/MFI |
| Polyoma Virus | 0/5 | - | - | 0/6 | - | - | 0/6 | RADIL | PCR/MFI |
| Clostridium piliforme | 0/5 | - | - | 0/6 | - | - | 0/6 | RADIL | PCR/MFI |
| Mouse Cytomegalovirus | 0/5 | - | - | 0/6 | - | - | 0/6 | RADIL | PCR/MFI |
| **BACTERIOLOGY (ANNUALLY)** | **6/3/2019** | | | **30/3/2020** | | | **7/4/2020** | | |
| Citrobacter rodentium | 0/5 | - | - | 0/6 | - | - | 0/6 | RADIL | PCR |
| Klebsiella oxytoca | 0/5 | - | - | 0/6 | - | - | 0/6 | RADIL | PCR |
| Klebsiella pneumoniae | 0/5 | - | - | 0/6 | - | - | 0/6 | RADIL | PCR |
| Proteus mirabilis | 0/5 | - | - | 0/6 | - | - | 0/6 | RADIL | PCR |
| Pseudomonas aeruginosa | 0/5 | - | - | 0/6 | - | - | 0/6 | RADIL | PCR |
| Salmonella sp. | 0/5 | - | - | 0/6 | - | - | 0/6 | RADIL | PCR |
| Bordetella bronchiseptica | 0/5 | - | - | 0/6 | - | - | 0/6 | RADIL | PCR |
| Corynebacterium kutscheri/bovis | 0/5 | - | - | 0/6 | - | - | 0/6 | RADIL | PCR |
| Pasteurella pneumotropica | 0/5 | - | - | 0/6 | - | - | 0/6 | RADIL | PCR |
| Pasteurella multocida | 0/5 | - | - | 0/6 | - | - | 0/6 | RADIL | PCR |
| Staphylococcus aureus | 0/5 | - | - | 0/6 | - | - | 0/6 | RADIL | PCR |
| Streptococcus pneumoniae | 0/5 | - | - | 0/6 | - | - | 0/6 | RADIL | PCR |
| Streptococcus sp. Beta hemolytic | 0/5 | - | - | 0/6 | - | - | 0/6 | RADIL | PCR |
| Streptobacillus moniliformis | 0/5 | - | - | 0/6 | - | - | 0/6 | RADIL | PCR |
| Helicobacter ganmani | 0/5 | - | | 0/6 | - | | 0/6 | | |
| Helicobacter hepaticus | 0/5 | - | - | 0/6 | - | - | 0/6 | RADIL | PCR |
| Helicobacter typhlonius | 0/5 | - | - | 0/6 | - | - | 0/6 | RADIL | PCR |
| Helicobacter mastomyrinus | 0/5 | - | - | 0/6 | - | - | 0/6 | RADIL | PCR |
| Helicobacter sp., H.bilis,H.rodentium,H.mastomyrinus | 0/5 | - | - | 0/6 | - | - | 0/6 | RADIL | PCR |
| **PARASITOLOGY (QUARTERLY)** | **6/3/2019** | **15/8/2019** | **15/12/2019** | **30/3/2020** | **28/9/2020** | **21/12/2020** | **7/4/2020** | | |
| Ectoparasites (e.g. fleas,fur mites, lice) | 0/5 | 0/5 | 0/5 | 0/6 | 0/6 | 0/6 | 0/6 | RADIL/SGH | PCR/ LM |
| Endoparasites (pinworms,tapeworms,roundworms) | 0/5 | 0/5 | 0/5 | 0/6 | 0/6 | 0/6 | 0/6 | RADIL/SGH | PCR/LM |

SingHealth

Defining Tomorrow's Medicine

Singapore Health Services Pte. Ltd
31 Third Hospital Avenue
#03-03 Bowyer Block C
Singapore 168753
www.singhealth.com.sg
UEN No. 200002698Z

Legends:
RADIL - Research Animal Diagnostic Laboratory
SGH - Singapore General Hospital
(SGH Department of Pathology)
MFI - Multiplex Fluorescent Immunoassay
IFA - Indirect Immunofluorescence Assay

**Supplementary Table 7: Specific pathogen free status of animal housing facility.** The SingHealth animal facility employs the sentinel method for surveillance of pathogens. Sentinel mice (ICR strain) are housed on soiled bedding removed from cages of other non-sentinel rodents in the population to be sampled. Sentinels are housed in individual ventilated cages like all other rodents. They are housed in the colony for at least 3 months before being tested. The table above details the specific pathogen tested for and the mode and frequency of testing. 0/6 indicates that zero out of six mice tested positive for that pathogen.
